# Disruption of Cholinergic Retinal Waves Alters Visual Cortex Development and Function

**DOI:** 10.1101/2024.04.05.588143

**Authors:** Timothy J Burbridge, Jacob M Ratliff, Deepanjali Dwivedi, Uma Vrudhula, Francisco Alvarado-Huerta, Lucas Sjulson, Leena Ali Ibrahim, Lucas Cheadle, Gordon Fishell, Renata Batista-Brito

## Abstract

Retinal waves represent an early form of patterned spontaneous neural activity in the visual system. These waves originate in the retina before eye-opening and propagate throughout the visual system, influencing the assembly and maturation of subcortical visual brain regions. However, because it is technically challenging to ablate retina-derived cortical waves without inducing compensatory activity, the role these waves play in the development of the visual cortex remains unclear. To address this question, we used targeted conditional genetics to disrupt cholinergic retinal waves and their propagation to select regions of primary visual cortex, which largely prevented compensatory patterned activity. We find that loss of cholinergic retinal waves without compensation impaired the molecular and synaptic maturation of excitatory neurons located in the input layers of visual cortex, as well as layer 1 interneurons. These perinatal molecular and synaptic deficits also relate to functional changes observed at later ages. We find that the loss of perinatal cholinergic retinal waves causes abnormal visual cortex retinotopy, mirroring changes in the retinotopic organization of gene expression, and additionally impairs the processing of visual information. We further show that retinal waves are necessary for higher order processing of sensory information by impacting the state-dependent activity of layer 1 interneurons, a neuronal type that shapes neocortical state-modulation, as well as for state-dependent gain modulation of visual responses of excitatory neurons. Together, these results demonstrate that a brief targeted perinatal disruption of patterned spontaneous activity alters early cortical gene expression as well as synaptic and physiological development, and compromises both fundamental and, notably, higher-order functions of visual cortex after eye-opening.

## Introduction

The maturation of nascent synaptic connections into highly ordered neural circuits is an integral feature of the developing nervous system that is regulated by the dynamic interplay between neuronal activity and genetic factors (Batista-Brito & Fishell 2009, Ben-Ari & Spitzer 2004, Fishell & Kepecs 2019, Stiles 2011, Weaver 2014). Previous work investigating how activity shapes the maturation of the primary visual cortex (V1) largely focused on sensory-dependent phases of visual circuit development. These studies showed that depriving animals of sensory experience after eye-opening led to lasting deficits in visual circuit wiring and visual function (Espinosa & Stryker 2012, Hensch 2005, Hensch & Fagiolini 2005, Reh et al 2020). However, many of the fundamental features of cortical circuit organization are established prior to the onset of eye-opening (Guillamon-Vivancos et al 2022, Jeon et al 2018, Li et al 2006, Martini et al 2021, Rochefort et al 2011). This suggests that innate genetic programs and pre-sensory forms of spontaneous activity, such as spontaneous retinal waves, might be relevant for cortical development and function.

Spontaneous neuronal activity is thought to be instructive for the assembly and maturation of multiple brain circuits across several species (Blankenship & Feller 2010). In the visual system spontaneous patterned bursts of action potentials within the retina, termed retinal waves, propagate across the developing visual system (Blankenship & Feller 2010). These cholinergic-mediated waves dominate visual system activity including the visual cortex within the first postnatal week in mice (∼P2–P8). This period is critical for the establishment of neocortical circuit connectivity and is a highly susceptible period for neurodevelopmental disorders (Allene & Cossart 2010, Danka Mohammed & Khalil 2020, Kroon et al 2019, Llorca & Deogracias 2022, Marin 2012, Wong & Marin 2019). Cholinergic retinal waves depend on the activation of nicotinic acetylcholine receptors containing the β2 subunit (β2-nAChRs) in retinal Starburst Amacrine Cells (Ford et al 2012, Ford & Feller 2012). Previous studies attempted to suppress cholinergic retinal waves in V1 through pharmacological suppression of β2-nAChR activity, enucleation, or genetic deletion of β2-nAChRs in β2−/− mice. However, those methods proved to be difficult to interpret (Bansal et al 2000, Blankenship & Feller 2010, Burbridge et al 2014, Cang et al 2005, Ford et al 2012, Ford & Feller 2012, Guil-lamon-Vivancos et al 2022, Kirkby et al 2013, Martini et al 2021, Stellwagen & Shatz 2002). Pharmacological suppression of β2-nAChRs is non-specific to the retina, and is often neurotoxic (Ackman et al 2012, O’Leary et al 1986, Penn et al 1998, Shatz & Stryker 1988, Sun et al 2008a); whereas enucleation or germline β2-nAChRs knockout (β2−/−) induces compensatory activity patterns in the retina, Superior Colliculus (SC), and V1 (Burbridge et al 2014, Cang et al 2005, Guillamon-Vivancos et al 2022, Sun et al 2008b). Moreover, the loss of β2 receptors in β2−/− mice has off-target effects that are not confined to the retina (Mazzaferro et al 2017) given that β2-nAChRs are also expressed in V1, as well as other various brain regions (Ackman et al 2012, O’Leary et al 1986, Penn et al 1998, Shatz & Stryker 1988, Sun et al 2008a). To overcome these problems, we restricted the removal of the β2 acetylcholine receptor subunit to a subregion of the retina using the retina-specific mouse line driver Pax6α-Cre in combination with the conditional mouse line β2fl/fl, thus generating Pax6α-Cre::β2fl/fl mice (β2-cKO mice). Because the Pax6α enhancer is selectively active in peripheral parts of the retina, β2-cKO mice reliably lack cholinergic waves in corresponding areas of the dorsal Lateral Geniculate Nucleus (dLGN) and SC but retain waves in the medial retinotopic region (Burbridge et al 2014). In contrast to β2−/− mice, β2-cKO mice do not have off-target effects because Pax6α is not expressed in the dLGN, SC or V1, leaving β2-nAChRs in these brain regions intact (Burbridge et al 2014, Marquardt et al 2001).

Here we show that disrupting cholinergic retinal waves disturbs typical gene expression programs in the postnatal V1 that are largely cell-type-specific. This results in the dysregulation of developmental transcriptomic profiles of multiple neuronal cell types, including layer 1 (L1) interneurons (IN) and pyramidal cells (PYR). Functionally L1 INs fail to establish normal connectivity and morphology during postnatal development. Furthermore, these postnatal synaptic changes in L1 INs and PYR are not recovered at later ages, with both cell types exhibiting developmentally arrested synaptic characteristics that persist past early postnatal development. These molecular and synaptic early alterations likely result in in vivo functional phenotypes. In particular, we show that cholinergic retinal waves are necessary for the development of V1 retinotopy. Higher order visual functions, such as the increase in visual response gain normally observed during periods of locomotion and arousal (active state) (Fu et al 2014, Niell & Stryker 2010, Vinck et al 2015) are also affected, with L1 INs and PYR having reduced state-dependent modulation of visual responses. Taken together, this work defines critical roles for pre-visual spontaneous activity in initiating a cascade of developmental molecular programs that shape the maturation and connectivity of cortical excitatory and circuits. These ultimately impact both basic visual properties, such as retinotopy, and higher order behaviorally relevant sensory processing, such as behavioral state modulation of visual activity.

## Results

### Selective removal of the β2 acetylcholine receptor subunit in subregions of the retina disrupts correlated spontaneous activity in the primary visual cortex

During the first week of life in mice, retinal waves are mediated by cholinergic neurotransmission through nicotinic acetylcholine receptors containing the β2 subunit (β2-nAChRs) (Ford et al 2012, Ford & Feller 2012). We adopted a conditional genetic strategy to selectively delete β2-nAChRs from restricted regions of the retina by crossing a β2fl/fl mouse line to the Pax6α-Cre driver line. Because the Pax6α enhancer is only expressed in two-thirds of the retina, specifically in the nasal and temporal regions (Fig. S1) (Marquardt et al 2001), β2fl/-::Pax6αCre (β2-cKO) mice exhibit reduced cholinergic waves in most but not all retinal regions (Burbridge et al 2014, Marquardt et al 2001). Because of the activity blockade in the retina, β2-cKO mice also lack cholinergic waves in regions of the SC that retinotopically correspond with the retinal regions that express Cre (Burbridge et al 2014). To test the impact of the loss of cholinergic waves in V1 resulting from this β2-cKO, we injected AAV9-CAG-GCaMP6f in V1 of P0 or P1 mice, and imaged wide-field calcium signals in awake animals at P7 (Fig. 1A), a time when retinal waves peak in wild type mice. As expected, we found that β2fl/-, Cre-negative littermates (control mice) exhibited retinal wave activity that travelled progressively through much of the surface of V1 (Fig. 1B), and lasted about 40-60 seconds with an inter-wave interval of about a minute (Fig. 1C, Fig. S1, Supp. Movie S1). In β2-cKO mice, the properties of these waves remained similar in the medial region of V1 (where cholinergic-mediated waves persists and which we denote as the wave+ region), which retinotopically corresponds to the Cre-negative territory of the retina (Fig. S1, Movie S2). However, the posterior part of the V1 (which we denote as the wave-region) showed a significantly decreased wave frequency (wave+ = 1.5+/-0.05 w/m, wave- = 0.83+/-0.07 w/m, Fig. 1C, Fig. S1, Supp. Video 2). In the wave-region relative to the wave+ region, wave size (wave+ = 3.1+/-0.8mm2, wave- = 1.2+/-0.1 w/m) and duration (wave+ = 16.6+/-3.2s, wave- = 4.2+/-0.8s) were both decreased while wave interval was increased (wave+ = 44.8+/-5.8s, wave- = 92.7+/-8.1s) (Fig. 1C). Notably, we did not observe large-scale compensatory activity reported in the less targeted wave knockout mouse models (Burbridge et al 2014, Cang et al 2008, Guillamon-Vivancos et al 2022, Sun et al 2008b). This indicates that our conditional genetic strategy is effective in manipulating spontaneous retinal activity in V1 during early postnatal development without inducing significant compensatory patterned activity.

**Figure 1:**
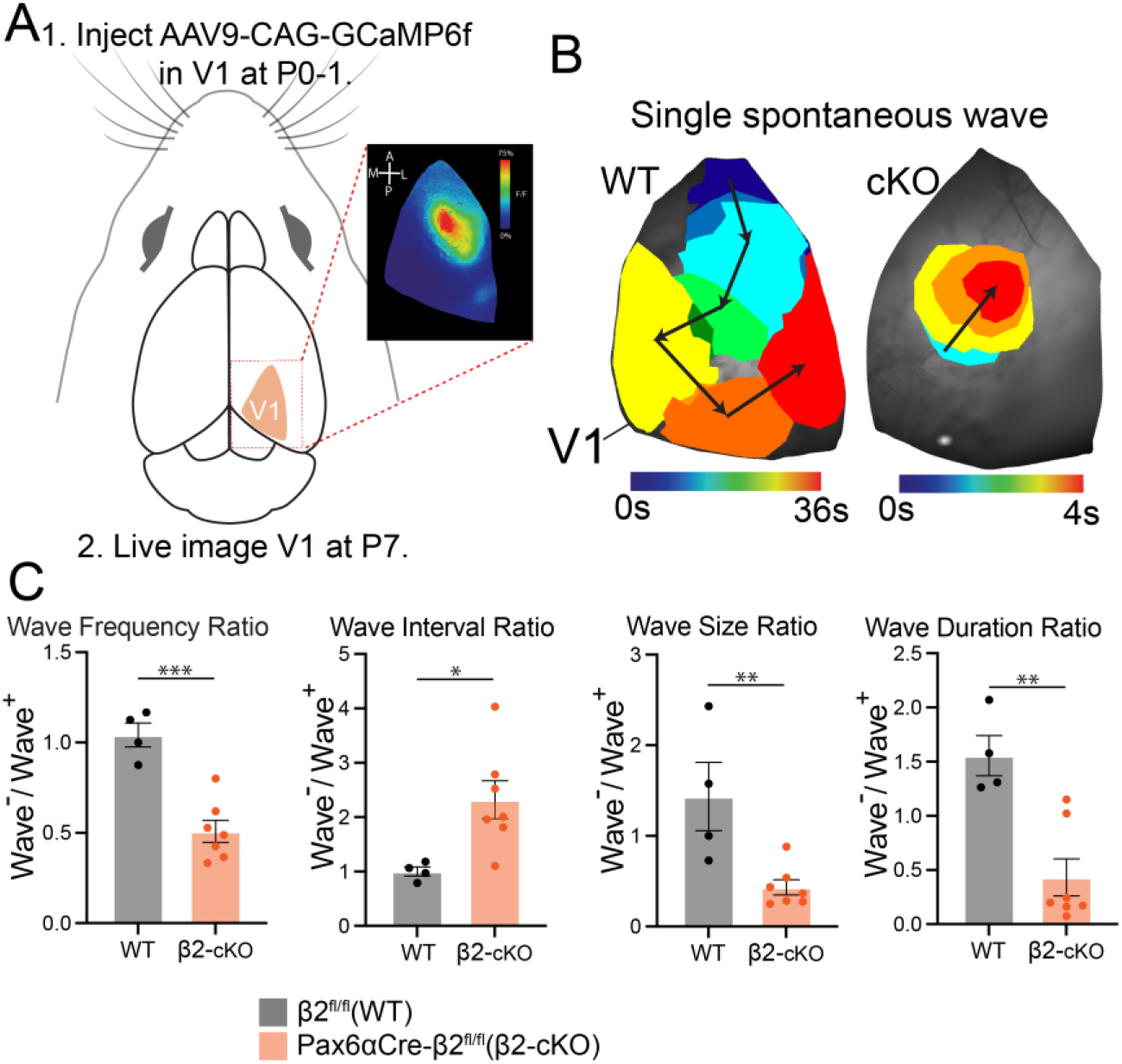
β2-cKO mutant mice have disrupted retinal waves in the visual cortex. A. Experimental setup. AAV9-CAG-GCaMP6f is injected at P0-1 and large-scale calcium imaging is done at P7 in V1 to characterize cortical spontaneous activity. Inset box shows an example of calcium wave activity in V1. B. Single spontaneous wave in cortex from WT (left) and β2-cKO (right) V1. WT waves generally last longer in duration and cover a larger percentage of V1 surface area than β2-cKO waves. C. Quantification of spatiotemporal properties of WT and β2-cKO V1 wave activity by region. Wave frequency (waves/minute) is lower in β2-cKO wave-region than in medial wave+ region with a corresponding increase in time between waves (Interwave Interval). Wave size (mm2) and duration (s) are smaller in β2-cKO wave-region. N=4 WT, 7 β2-cKO. *** p<0.001, ** p<0.01, * p<0.05.

### Retinal wave disruption alters gene expression programs in developing excitatory and inhibitory neurons in V1

To determine whether retinal wave disruption impacts cortical development at the molecular level, we profiled gene expression across medial (wave+) and posterior (wave-) regions of V1 in β2-cKO mice using single-nucleus RNA-sequencing (snRNAseq) at P7, when cholinergic retinal waves are prevalent. As a control, we also performed snRNAseq on medial and posterior V1 tissue from Cre-negative β2fl/-littermates (control mice). Wave+ and wave-regions of V1 in β2-cKO mice were identified by wide-field imaging of Ca2+ dynamics via panneuronal expression of AAV9-CAG-GCaMP6f (Fig. 1), after which each region was micro-dissected and flash-frozen for later processing (see Methods). Nuclei were isolated and sequenced across three biological replicates using the inDrops strategy of high-throughput transcriptomic profiling (Zilionis et al 2017), then subjected to quality control processing followed by analysis via Seurat and Monocle (Qiu et al 2017, Stuart et al 2019). In total, 65,744 nuclei were included in the final dataset, the majority of which belonged to neurons (Fig. 2A-D; Fig. S2A).

**Figure 2.**
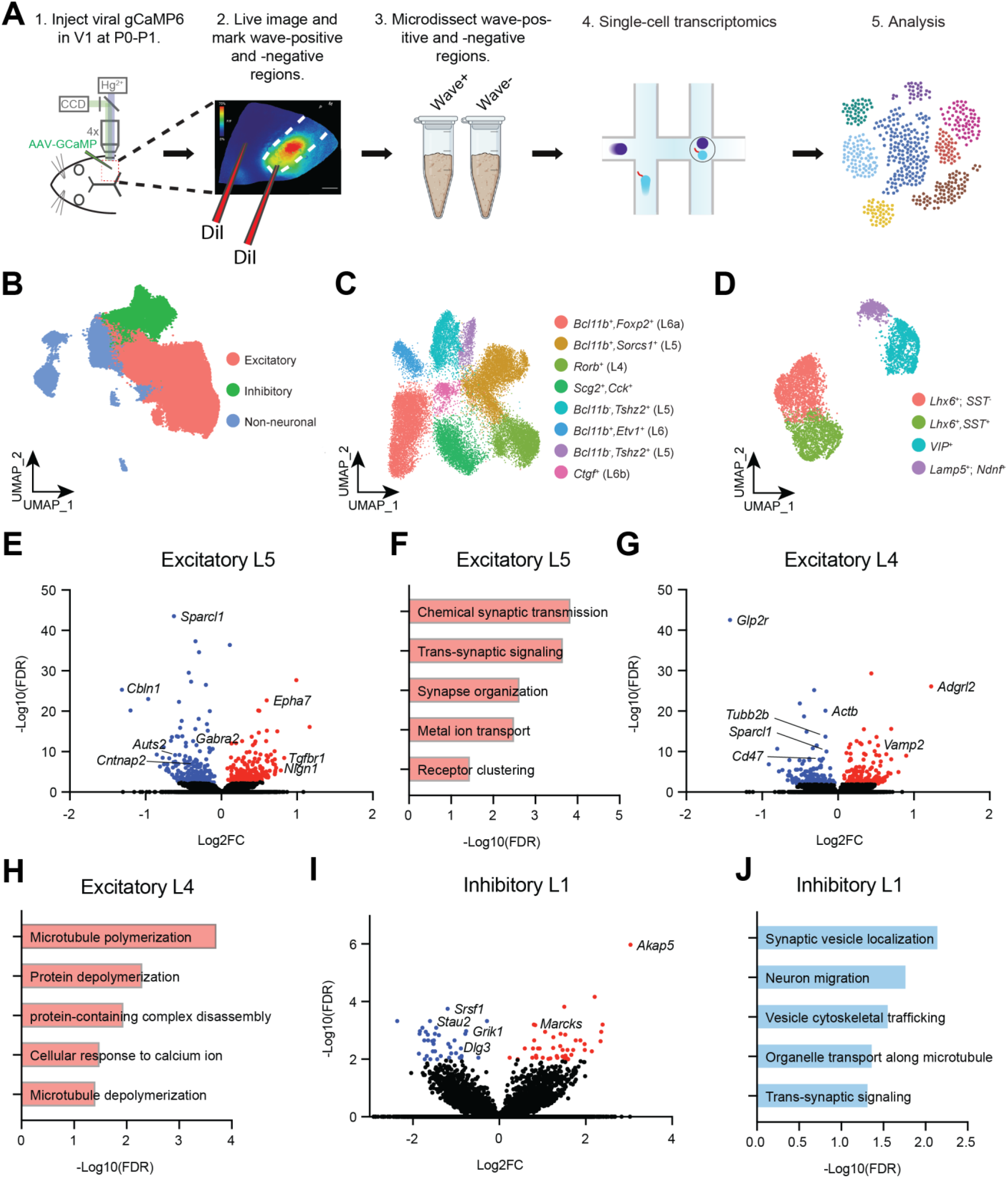
Wave disruption modifies the transcriptomic profiles of developing cortical cells. (A) Schematic describing the identification, isolation, and sequencing of wave+ (medial) and wave-(posterior) regions of V1 in β2-cKO mice. (B) Two-dimensional UMAP (Uniform Manifold Approximation and Projection for Dimension Reduction) representation of all cell types in the dataset by general cell class. (C) UMAP of excitatory cell types in deep cortical layers (L4 – L6). (D) UMAP of inhibitory cell types across all layers. (E),(G),(I) Volcano plots demonstrating transcripts that are more highly (red) or more lowly (blue) expressed in layer 5 excitatory neurons (E), layer 4 excitatory Rorb+ neurons (G), and layer 1 Ndnf+ inhibitory neurons (I) in wave+ versus wave-V1 of β2-cKO mice. (F),(H),(J) Gene ontology categories enriched among genes that are more highly expressed in wave+ versus wave-V1: L5 excitatory neurons (F), L4 excitatory neurons (H), and L1 inhibitory neurons (J).

Applying differential gene expression analysis, significant differences in gene expression were observed between medial and posterior V1 regions across all neuronal types in both β2-cKO and control mice (Fig. S2B-C). The observation of region-specific gene expression in mice with normal retinal waves is consistent with the presence of transcripts that may be organized retinotopically. Our data suggest that some features of region-specific gene expression observed in control mice are preserved in β2-cKO mice while others are not. For example, the synaptogenic adhesion molecule Cadm1 is expressed much more highly in posterior V1 compared to medial V1 in both β2-cKO and control mice (Fig. S3). In contrast, transcripts encoding other known synaptic organizers, such as Sema3a and EphB2, lose their regional specificity when waves are disrupted. Thus, one potential role for retinal waves may be to aid in the establishment of molecular organization across the retinotopic axis in V1.

To identify retinal wave-dependent changes in gene expression, we next focused our analysis on the transcripts that were differentially expressed between medial (wave+) and posterior (wave-) regions of V1 in β2-cKO. We observed that differentially expressed genes tended to be cell-type-specific, consistent with prior data indicating that activity-dependent gene expression in neurons often differs between neuronal subtypes (Hrvatin et al 2018) (Fig. S3). Given that retinal waves coincide temporally with the assembly and refinement of thalamocortical circuitry, we hypothesized that retinal wave blockade disrupts the expression of some genes that encode proteins involved in synapse development, remodeling, and function. Consistent with this hypothesis, genes that are downregulated as a result of retinal wave blockade in Rorb+ excitatory neurons of L4 and pyramidal neurons of L5 and L6 included mediators of synaptic organization and assembly, such as the postsynaptic scaffolding molecule Shank3 and the synaptogenic adhesion molecule Neuroligin 1, both of which are autism risk genes (Fig. 2E-H) (Monteiro & Feng 2017, Trobiani et al 2020). Finally, given that the integration of inhibitory neurons into cortical circuits also temporally coincides with cholinergic waves, we examined the transcriptional differences in inhibitory neurons between wave+ and wave-regions in β2-cKO mice. Among the four inhibitory classes analyzed, changes in the expression of synaptic genes were most prevalent in layer 1 interneurons that express Ndnf (NDNF-INs), and in parvalbumin interneurons (PV-INs) which predominantly reside in recipient layer 4 (Fig. 2I, J). These results are consistent with interneurons in both L1 and L4 directly receiving perinatal thalamic inputs, indicating that these neuronal classes reside in regions of cortex that are likely to be highly susceptible to the influence of early retinal activity on circuit organization. Overall, these data are consistent with retinal wave activity playing an important role in shaping the molecular development of V1 postnatally

### Retinal wave disruption impairs synaptic development of pyramidal cells

During the first postnatal weeks neocortical neurons receive exuberant synaptic connectivity that is later pruned in an activity-dependent manner (Antonini et al 1999, Antonini & Stryker 1996, Faust et al 2021). Much of our understanding of early pruning and refinement events comes from the retinogeniculate system in which retinal waves are critical for this refinement, and from the refinement of V1 connectivity after eye-opening. Here we test the hypothesis that, in the absence of activity driven by retinal waves specifically, PYR in the V1 might inappropriately retain an increased number of synaptic inputs as a result of the initial over-production of synapses and/or impairments in activity-dependent synaptic pruning. This hypothesis is consistent with our observation that pyramidal neurons exhibit early alterations in levels of synapse-associated genes when waves are blocked (Fig. 2E-H). We performed in vitro electrophysiology of PYR in acute slices from posterior regions (wave-) of V1 in β2-cKO and control animals between P25 and P30 (Fig. 3A). Miniature excitatory postsynaptic currents (mEPSCs), a functional measure of synaptic strength and synapse number, were recorded at a holding potential of -70mV in the presence of tetrodotoxin and GABA receptor antagonists (Fig. 3B). Our data shows that disruption of retinal waves leads to an increase in excitatory neurotransmission onto neocortical PYRs located in the thalamocortical recipient layers (layers 4 and 5) in β2-cKO mice, as evidenced by increased EPSC rate and amplitude (Fig. 3B-C), and increased number of excitatory synapses containing VGlut1 (Fig. 3C), suggesting that retinal waves may be necessary for the elimination of excess excitatory inputs to V1 (Fig. 3C-E, Fig. S4). In line with this possibility, L4 excitatory neurons in wave-regions of V1 in β2-cKO mice exhibit heightened early expression of CD47 (Fig. 2G), a signaling protein that inhibits synaptic pruning by microglia (Lehrman et al 2018). To evaluate if inhibition is altered in β2-cKO mice, miniature inhibitory postsynaptic currents (mIPSCs) were recorded at 0 mV in the presence of tetrodotoxin and antagonists of AMPA and NMDA receptors. Our data shows that β2-cKO mice have reduced mIPSC amplitude (Fig. 3E-F), and reduced number of L4 PV synapses (Fig. 3G), suggesting that inhibitory neurotransmission onto PYR is also impaired. Taken together, these observations suggest that β2-cKO mice exhibit abnormal synaptic connectivity during both early (P8) and later (P25 – P30) stages of development that are most prevalent in the V1 input layers, and are likely to lead to impaired E/I balance

**Figure 3:**
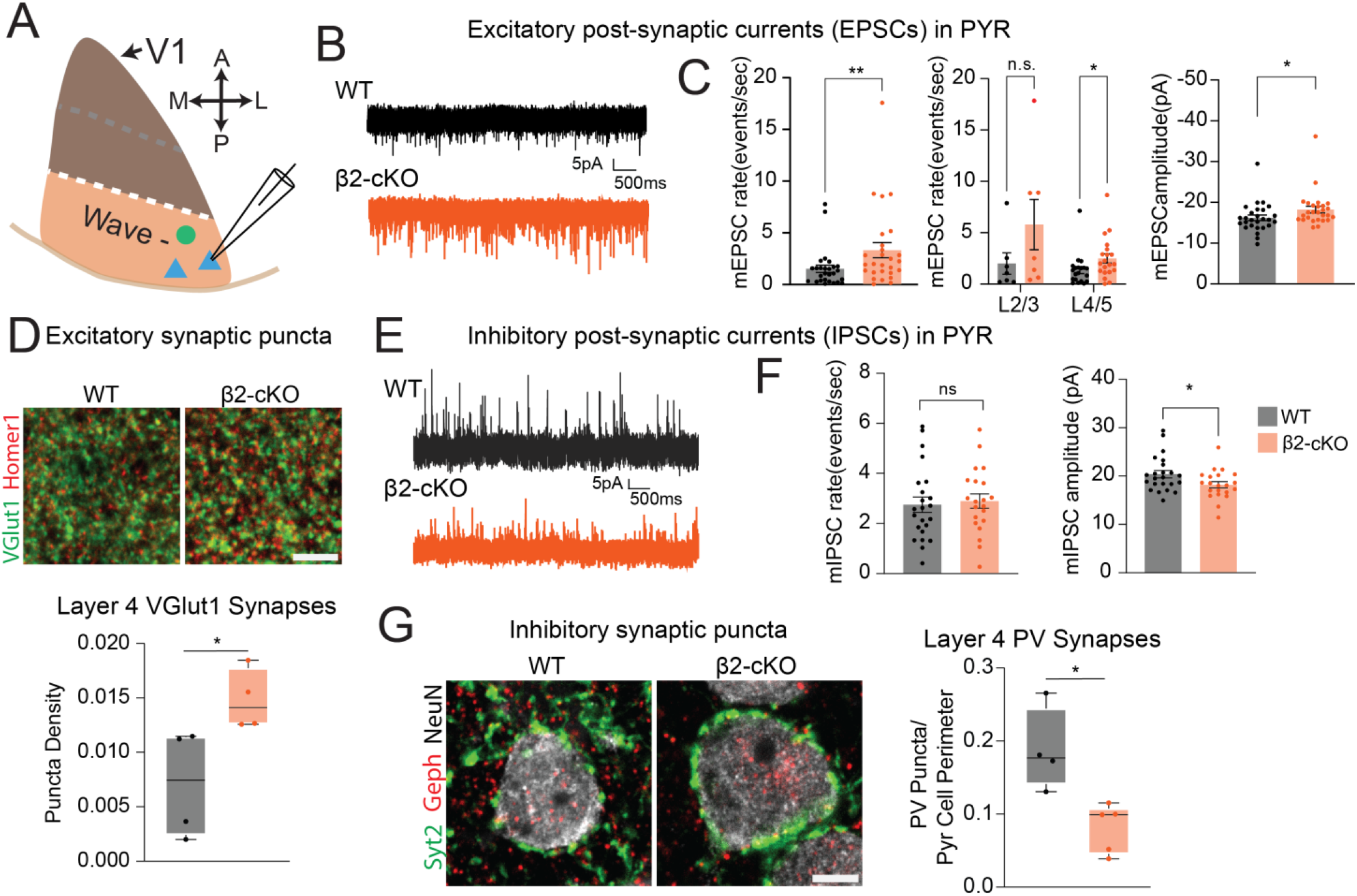
V1 pyramidal cells (PYR) in β2-cKO mutant mice have increased excitatory inputs and reduced inhibitory inputs. A. Schematic of experimental approach. PYR were randomly patched in posterior wave-regions of the V1 to record mEPSCs and mIPSCs in control and mutant animals. B. Representative traces of excitatory post-synaptic currents onto pyramidal cells. C. Average frequency of mEPSCs in control and β2-cKO pyramidal cells across all cortical layers, in superficial and deep layers, and average amplitude of mEPSCs in control and β2-cKO pyramidal cells across all cortical layers. D. Top, representative images of excitatory synaptic puncta in layer 4 of P21 animals. Bottom, quantification of puncta density. E. Representative traces of inhibitory post-synaptic currents onto pyramidal cells. F. Average frequency and amplitude of mIPSCs in control and β2-cKO pyramidal cells across all cortical layers. G. Left, representative images of inhibitory synaptic puncta in layer 4 of P21 animals. Right, quantification of puncta density. N=6 control and mutants for physiology, N=4-5 control and mutants for synapse analysis.

### Retinal waves are necessary for the development of V1 retinotopy

Given that retinotopic organization in mouse V1 initiates at neonatal ages before the onset of vision at eye-opening (Blankenship & Feller 2010, Huberman et al 2008, Kirkby et al 2013), we sought to determine whether cholinergic retinal waves are required for V1 retinotopy. We performed in vivo extracellular recordings in awake, head-fixed β2-cKO animals and littermate controls between P25 and P30 (Fig. 4A), a time frame by which visual tuning has already been established (Jeon et al 2018, Li et al 2006, Rochefort et al 2011). We used 64 channel linear silicon probes to record the firing activity of regular spiking (RS) cells (putative excitatory neurons), and fast spiking (FS) cells (putative PV-INs) across all cortical layers targeting the posterior lateral (wave-) region of V1 (Fig. 4A). We recorded 697 well-isolated single units (356 in control [298 regular spiking and 58 fast spiking], and 341 in β2-cKO [280 regular spiking and 61 fast spiking] in β2-cKO). Mice were shown visual stimuli contralateral to the recording site consisting of small (20° visual angle), circular, drifting sinusoidal gratings of varying orientations, temporal frequency, spatial frequency, size, and elevation. To examine retinotopy, drifting grating stimuli were shown at multiple elevations between -20° to 45° (with 0° being horizontally in line with the mouse eyes). Previous work has shown that the neurons in the lateral, posterior V1 (our targeted region), prefer stimuli at higher elevations, and that neurons in a given spatial area are likely to have similar retinotopic tuning preferences (Cang et al 2005, Marshel et al 2011). Our results show that suppression of retinal waves in β2-cKO mice disrupted retinotopy, particularly compromising elevation maps (Fig. 4C-E), while sparing other receptive field features (Fig. S5). Population analysis indicated that control animals have narrow average (or similar) (Fig. 4C, D), responses that peaked at 40°(high elevation) (Fig. 4C), consistent with the retinotopic preference of lateral posterior V1 (Marshel et al 2011). In contrast the average responses of β2-cKO animals were broader (less similar) (Fig. 4C), and peaked at lower elevation, around 20° (Fig. 4C). To analyze the preferred elevation of individual neurons, single unit responses to stimuli at different elevations were fit with gaussian tuning curves (Fig. 4D, Fig. S5). Analysis of the preferred elevation of individual neurons showed a similar effect to population level responses, with individual neurons from control mice preferring elevations clustered near 40° (39.7 ±9.0 s.d., N=5/6 mice with units visually responsive to elevation stimuli), while neurons from β2-cKO mice spread across elevations (Fig 4D; 22.0 ±13.9 s.d., N=5/6 mice with units visually responsive to elevation stimuli). We found no difference in RS vs FS cells and as such present them together. The increased variability in elevation preference in β2-cKO mice compared to the tightly clustered preferred elevations in control mice indicate disrupted retinotopic organization of visual responses. This is consistent with snRNAseq data suggesting the partial disruption of region-specific gene expression organized across the retinotopic axis in wave-vs wave+ regions of β2-cKO mice at P7 (Fig. 2). We further examined the variability of preferred locations of neurons within single animals and determined that β2-cKO animals have greater within animal variability suggesting that the difference in elevation tuning is not due to recording site variation across β2-cKO animals that did not occur in control mice (Fig 4E). These data demonstrate that spontaneous, patterned retinal waves during the first week of life contribute to the establishment of retinotopy in the V1.

**Figure 4:**
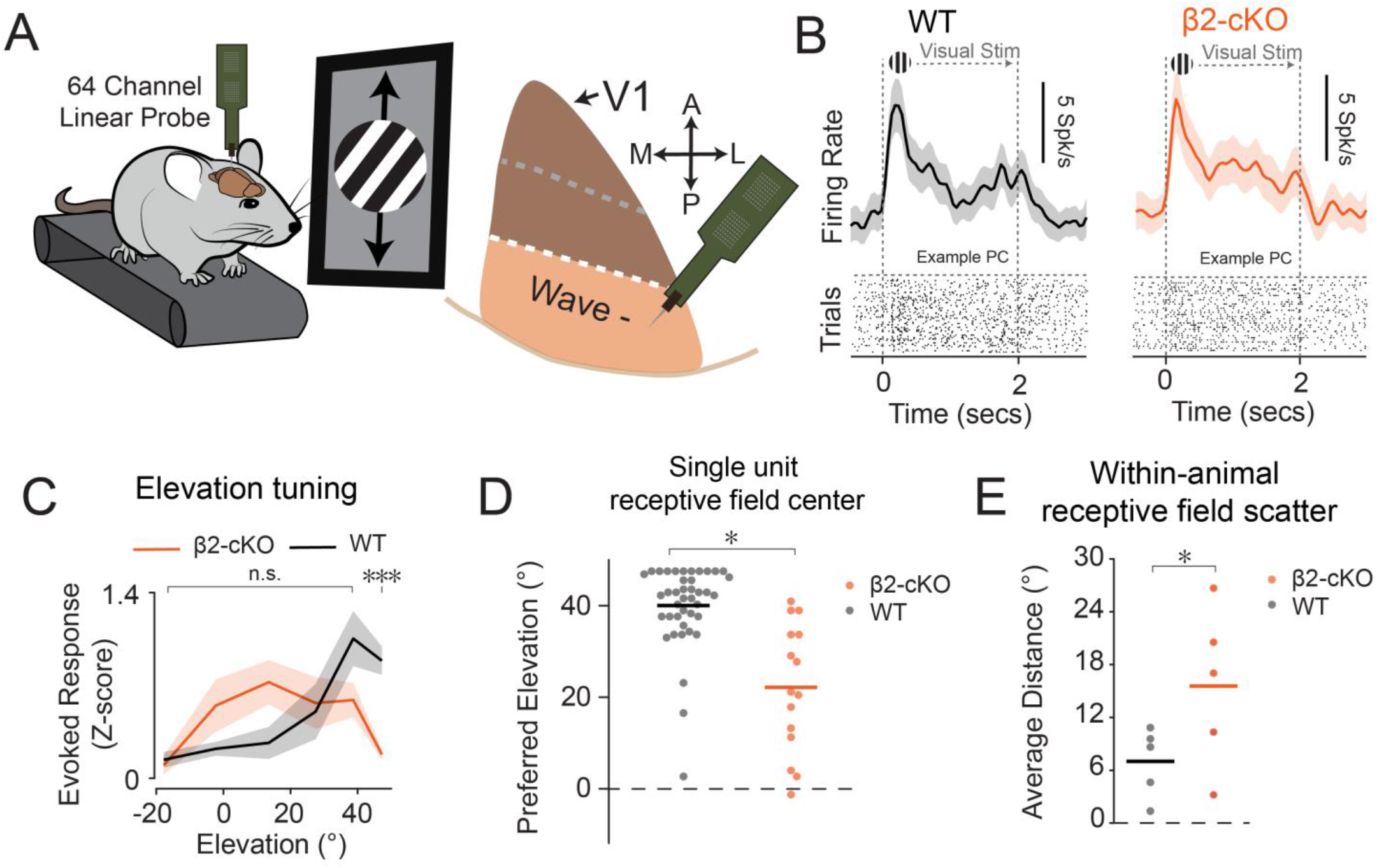
β2-cKO mutant mice have compromised retinotopy in the visual cortex. A. Experimental schematic showing head fixed mice with electrophysiological recording during visual stimulation and target area in wave negative posterior visual cortex. B. Example units from both control (Pax6α-Cre; black) and Pax6α:β2-cKO (Beta2fl/fl::Pax6α-Cre; red) animals. Top: mean firing rate for visual response to preferred stimulus. Bottom: Raster plots of spiking activity for stimulus from above. C. Animal grand average z-scored spiking response to varied elevation of visual stimulation (Two-sample t-test, p < 0.001). D Preferred elevation of individual units determined as peak of gaussian tuning curve fit, bar representing mean (p < 0.001, Mann Whitney U test). N=5 controls and 5 mutants, n = 41 and 15 units, respectively. E. Average pairwise distance of receptive field centers within mice p = 0.03, one-tailed Student’s t-test. N=5 controls and 5 mutants.

### Retinal wave disruption impairs synaptic maturation of layer 1 cortical interneurons

The integration of inhibitory interneurons (INs) into cortical circuits occurs during the first week of life, and L1 IN connectivity in particular is likely to be impacted by spontaneous activity due to the presence of strong thalamocortical projections in L1 at this stage (Ibrahim et al 2021). Additionally, since L1 INs play a crucial role in shaping sensory maps (Che et al 2018, Takesian et al 2018), the disrupted retinotopic map observed in β2-cKO mice lead us to hypothesize that the postnatal development of L1 INs might rely on cholinergic retinal waves. Accordingly, our transcriptomic analysis indicates that the synaptic development of layer 1 INs may be impaired when retinal waves are blocked (Fig. 2I-J). To directly test if cholinergic retinal waves impact the synaptic maturation of layer 1 INs, we measured excitatory synaptic connectivity onto layer 1 INs in acute brain slices from control and β2-cKO mice (Fig. 5A). Measurements were done around the end of the second postnatal week, a period during which layer 1 INs synaptic maturation is occurring, and layer 1 INs shift from predominantly receiving thalamic connectivity to establishing other long-range connectivity including cholinergic connectivity (Ibrahim et al 2021). Layer 1 INs showed a reduction in mEPSC rate and increase in mEPSC amplitude in β2-cKO compared to control mice, indicating that these cells receive altered excitatory inputs in the absence of retinal waves (Fig. 5B-C). Given that both glutamatergic and cholinergic inputs contribute to EPSCs, we isolated the contribution of cholinergic inputs toward the excitation by filtering the mEPSCs for slower kinetics (Fig. S6, see methods for details), which revealed that the rate of cholinergic inputs was significantly reduced in β2-cKOs (Fig. 5D), suggesting that L1 INs have disrupted cholinergic inputs from the basal forebrain (Ibrahim et al 2021). Morphologic tracing of the recorded cells demonstrated that total dendrite length was unchanged while axon length was significantly reduced in β2-cKO group (WT axon length = 4390 +/-1025 μm, β2-cKO axon length = 1471 +/-480 μm, Fig. 5E-F), implying that both the excitatory input as well as inhibitory output of L1 INs are influenced by patterned spontaneous activity. Overall, this disruption suggests that L1 INs may be impaired in shaping sensory maps during development (Che et al 2018, Takesian et al 2018) and may have disrupted in vivo activity patterns and connectivity.

**Figure 5:**
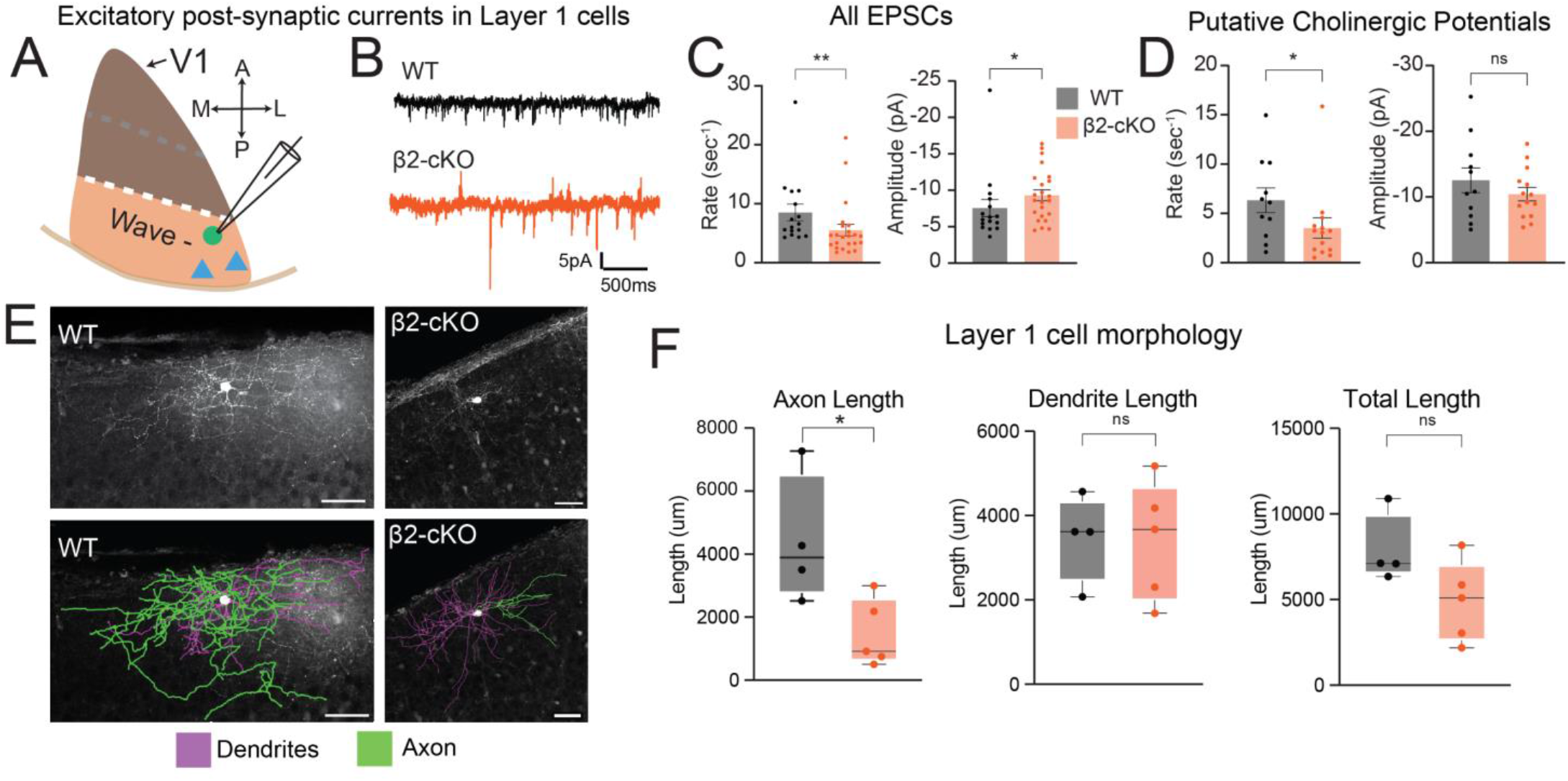
Layer 1 interneurons in β2-cKO mutant mice have altered excitatory inputs. A. Schematic of experimental approach. Cells in layer 1 were patched in posterior, wave-negative V1. B. EPSCs onto layer 1 cells in control and β2-cKO mutant mice. C. have mEPSCs have increased amplitude but a decreased rate (n = 16 WT and 23 cKO, p = 0.004 rate, p = 0.034 amplitude, Mann-Whitney). D. Putative cholinergic potentials exhibit a decreased rate but no change in amplitude (n = 11 WT and 14 cKO p = 0.038 rate, p = 0.467 amplitude, insert stats, Mann-Whitney). E. Filled layer 1 neurons were traced to determine gross morphology. Scale = 50□ m F. Total length of axons plus dendrites and dendrite length alone is not significantly different between groups, but axon length is significantly shorter in mutants. N=4 control and 3 mutants (5 mutants for morphology). * p<0.05; **p<0.01

### Retinal wave disruption impairs the behavioral state related activity of layer 1 cortical interneurons

We next hypothesized that the altered cholinergic transmission on L1 INs might disrupt their activity during behavior states modulated by acetylcholine, and sought to test how the absence of retinal waves impacts the in vivo activity of layer 1 INs and influences the behavior state modulation of this neurons. We used 2-photon calcium imaging to measure the in vivo spontaneous and visually evoked activity of layer 1 INs. We injected neonatal β2-cKO and control mice with an AAV-Dlx6-GCaMP6f vector to express the calcium indicator in inhibitory neurons in the visual cortex (Fig. 6A). We implanted these mice with cranial windows and imaged activity of layer 1 INs between P25 and P30 (Fig. 6B,C), when tuning of neurons in the visual cortex have achieved maturity (Jeon et al 2018, Li et al 2006, Rochefort et al 2011). Previous work has shown that Layer 1 IN activity is highly modulated by the behavioral state of the animal (Bugeon et al 2022, Cohen-Kashi Malina et al 2021). We therefore looked at the behavioral state modulation of layer 1 INs by comparing their activity across changing arousal states of the animal. As expected, layer 1 INs in control animals are highly modulated by behavioral state in the absence of visual stimulation (Fig. 6D). In comparison, β2-cKO animals showed reduced arousal evoked activity compared to controls (Fig. 6D, E), consistent with the observation that cholinergic neuromodulation onto layer 1 INs is reduced in mutant animals (Fig. 5D). To test if the functional connectivity onto layer 1 INs is compromised in β2-cKO mice we measured their visually evoked responses. As expected, layer 1 INs are visually responsive (Fig. 6F), however the magnitude of the visual response is increased in β2-cKO mice (Fig. S7), consistent with the idea that neurons in β2-cKO mice do not mature properly and receive stronger thalamic connectivity than controls. In the V1, the magnitude of sensory responses is highly influenced by the animals’ behavioral state (McGinley et al 2015, Stringer et al 2019), and previous work has shown that sensory-evoked activity of L1 INs is strongly enhanced when an animal is aroused (Bugeon et al 2022, Cohen-Kashi Malina et al 2021). We therefore analyzed the behavioral-state modulation of visual responses in layer 1 INs. To compare the behavioral state-dependent visually evoked responses, we quantified the baseline corrected visually evoked signal in controls and β2-cKO animals. We found that β2-cKO mice have reduced state modulation of visual responses (Fig. 6F, G), indicating that disruption of cholinergic retinal waves results in state-dependent dysregulation of layer 1 INs likely in part by altering cholinergic connectivity onto layer 1 of V1 (Fig. 5D).

**Figure 6:**
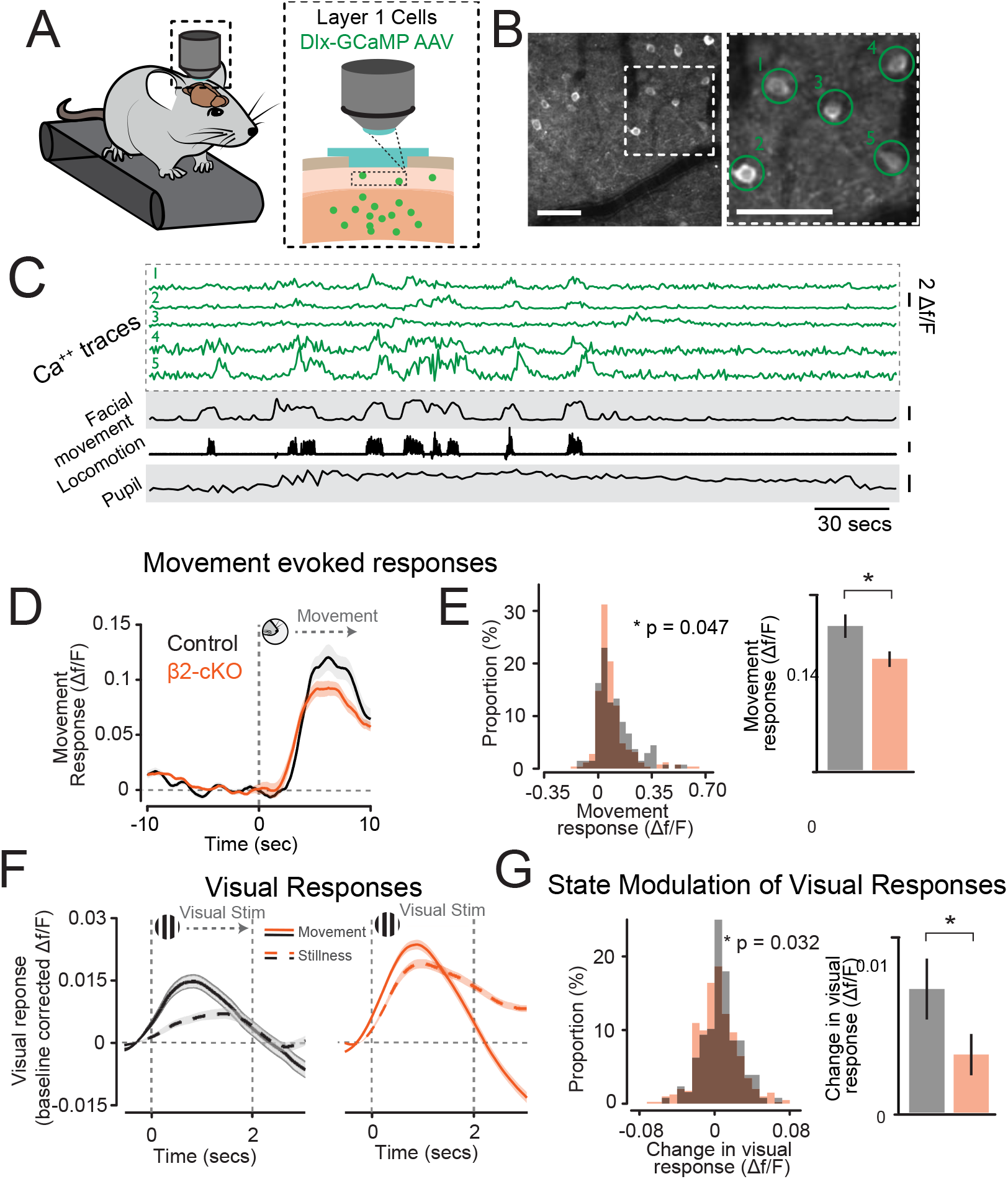
Layer 1 interneurons in β2-cKO mice exhibit reduced modulation by behavioral state. A. Schematic of the imaging scheme from layer 1 INs. B. Example field of view from imaging experiments. (Scale bar 100um) C. Example traces showing simultaneous recordings of Ca signals from Layer 1 interneurons with measurements of state including facial movement, treadmill locomotion, and pupillometry. Scale bars represent 50% of maximum value for facial movement and pupil and 10 cm/s for locomotion. D. Peri-movement average of cell activity locked to the beginning of facial movement bouts. E. ß2-cKO mice exhibit decreased movement responses (p = 0.047, Mann-Whitney U test) Left: histogram of all cells. Right: population mean and s.e.m. F. Peristimulus average of cell activity locked to visual presentation and normalized to prestimulus activity level (df/FSTIM – df/FITI) divided by state and by genotype. Left: WT, right: Pax6α:β2-cKO. G. ß2-cKO mice exhibit decreased difference across state in average visually evoked df/F values in each cell recorded. (p = 0.032, Mann-Whitney test) Left: histogram of all cells. Right: population mean and s.e.m. N = 8 animals, n = 385 cells.

### Retinal wave disruption reduces state-dependent visual activity of pyramidal cells

Having established that retinal waves are required for the state modulation of layer 1 INs, and given that arousal associated L1 INs gain modulate local excitatory neurons (Cohen-Kashi Malina et al 2021), we next sought to determine whether cholinergic retinal waves contribute to the behavioral state-dependent modulation of excitatory neurons. We examined neuronal spiking activity during visual stimulation according to behavioral state by dividing visual trials based on extent of movement (Fig. 7A-C). Our data show that β2-cKO mice display robust visually driven spiking responses (Fig. 7B, C), here calculated as signal-to-noise-ratio or SNR ([FRevoked – FRbaseline] / FRbaseline), as well as changes in spontaneous firing rates and oscillatory activity with changes in arousal state (Fig. S8). However, when we separated visual trials based on behavioral state, we observed differences in the magnitude of visual responses between control and β2-cKO mice specifically during trials when mice displayed high levels of movement (Fig. 7B-E). As previously reported, control animals showed increased visual responsiveness during periods of high arousal compared to periods of low arousal (Fig. 7B-E). However, visual responses in β2-cKO animals were similar regardless of an animal’s behavior, indicating a lack of state-dependent modulation of visual responses when retinal waves were disrupted (Fig. 3B-E). Taken together, our data show that disruption of cortical spontaneous waves during postnatal development not only impacts retinotopy at later ages but also impairs higher order visual responses such as gain modulation by arousal state. These data implicate retinal waves specificallyin the development of state-dependent visual function.

**Figure 7:**
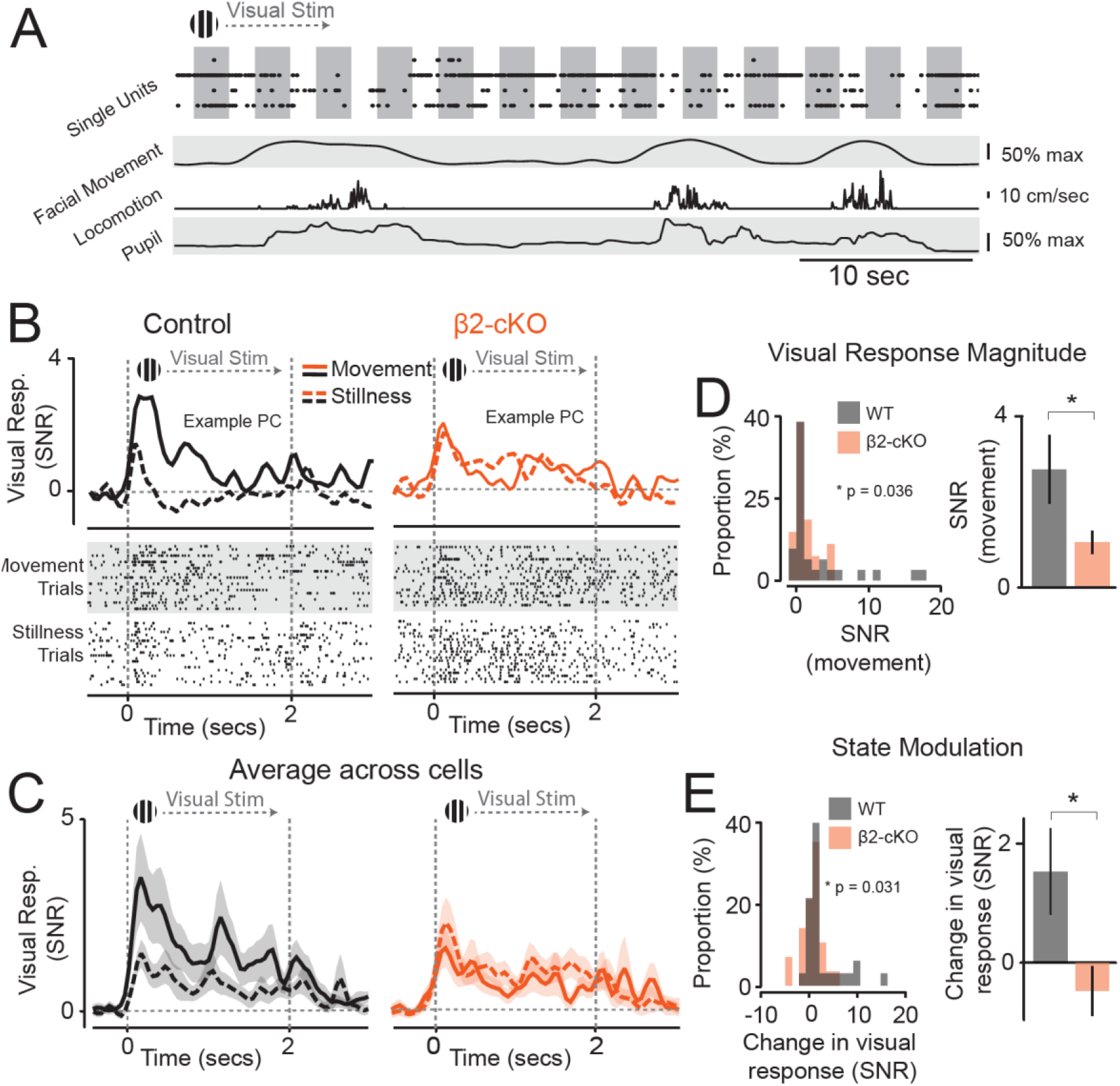
Pax6α-Cre::Beta2fl/fl mice exhibit reduced behavioral-state modulation of visual responses. A. Example data showing simultaneous recording of spiking activity and LFPs with measure of behavioral state (locomotion, facial movement, and pupil diameter). Scale: 50% max facial movement and pupil, 10cm/s locomotion. B. SNRs in example regular spiking cells (RS) from control (Pax6α-Cre; black) and mutant (Beta2fl/fl::Pax6α-Cre; red) animals across behavioral states (stillness vs. movement). Note reduced modulation of behavioral state in the mutant. Top: mean SNR for visual response to preferred stimulus. Bottom: Raster plots of spiking activity for stimulus from above. C. Population signal to noise ratio (SNR) for visual responses during movement and stillness calculated across units for control (black, n=31) and mutant (red, n=27) animals. D. ß2-cKO mutants exhibit decreased average SNR during first 500ms of stimulation. (p = 0.036, Welch’s t-test). Left: histogram of all units. Right: population mean and s.e.m. E. ß2-cKO mutants exhibit decreased change in SNR from stillness to movement (p=0.03 Welch’s t-test) Left: histogram of all units. Right: population mean and s.e.m. N=5 controls and 5 mutants.

## Discussion

Our results show that cholinergic retinal waves are necessary for maturation and connectivity of both inhibitory and excitatory neurons in the visual cortex, and that their absence causes long lasting impairments to first order visual function, such as retinotopy, and also impairs higher order visual processing, such as state-dependent sensory gain.

The roles of spontaneously generated activity, such as retinal waves, in the establishment and function of sensory cortical circuits have, in the past, been difficult to investigate largely due to a lack of tools for precisely manipulating spontaneous patterned activity during relevant phases of development. Previous manipulations of cholinergic retinal waves using a global β2 knockout (β2-/-) mouse line (Bansal et al 2000) suggested that cholinergic retinal waves may be relevant for retinotopy and eye-specific segregation (Cang et al 2005, Chandrasekaran et al 2005, Grubb & Thompson 2004, McLaughlin et al 2003, Mrsic-Flogel et al 2005, Shah & Crair 2008, Stafford et al 2009, Triplett et al 2009, Wang et al 2009). However, subsequent reports revealed that retinal waves are still present, and p¬¬ossibly even more frequent, in β2−/− mice compared to wild-type controls, and identified the presence of compensatory activity in brain regions down-stream of the retina in β2−/− mice (Stafford et al 2009, Sun et al 2008a, Sun et al 2008b). Due to these caveats, it remains unclear whether cholinergic retinal waves coordinate the development and function of visual cortex. Here, we have overcome these challenges by using a conditional genetic strategy whereby we selectively remove β2 subunits from part of the retina. We show that the β2-cKO mouse model largely avoids patterned, compensatory activity allowing us to assess the specific effects of cholinergic retinal wave disruption on cortical development.

Sensory experience and activity-dependent gene expression are critical for multiple stages of synapse and circuit development (Yap & Greenberg 2018). However, while activity-dependent gene programs induced by visual experience have been previously described (Cheadle et al 2018, Hrvatin et al 2018), the control of transcription in V1 by retinal waves has remained largely unexplored. Our results demonstrate that spontaneous activity generated in the retina coordinates gene programs in V1 that are largely cell-type-specific. The snRNAseq datasets presented here provide transcriptome-wide atlases of retinotopically organized and spontaneous activity-dependent gene programs in V1 during postnatal development. The perinatal changes in excitatory and inhibitory neurons in our dataset contains clues regarding the molecular factors that allow retinal waves to influence the establishment of connectivity. Of particular relevance, our data shows that cholinergic retinal waves impact the synaptic molecular machinery of layer 1 INs and pyramidal cells in the V1 input layers in early development. We find that these changes correspond with altered synaptic connectivity onto layer 1 INs, and pyramidal cells in the V1 input layers.

The elaboration of nascent synaptic connections into highly ordered neural circuits is an integral feature of the developing nervous system (Ascoli et al 2008, Wong & Marin 2019). Synaptic maturation during neonatal development is a highly dynamic and precisely timed process that involves various feedback loops between different cell types, and is likely to be impacted by spontaneous activity (Faust et al 2021). Cortical synapses are overproduced early in postnatal development, after which connectivity is pruned in an activity dependent manner (Batista-Brito & Fishell 2009, Belousov & Fontes 2013, De Marco Garcia et al 2011, Fukuda & Kosaka 2000a, Fukuda & Kosaka 2000b, Galarreta & Hestrin 2002, Ibrahim et al 2021, Pereda 2016, Pouchelon et al 2021, Sohl et al 2005, Tuncdemir et al 2016, Zolnik & Connors 2016). As shown here, this process is impacted by cholinergic retinal waves, as these represent a prevalent source of neural activity during early cortical postnatal development. Accordingly, we observed that PYR located in the thalamic input cortical layers (such as layer 4) of β2-cKO mice have increased excitatory transmission. In addition to changes of connectivity onto PYR in the input layers, we also observed that synaptic transmission onto layer 1 INs was altered. Layer 1 INs receive strong thalamic input around P7, after which they transition towards receiving long range connectivity from other regions (De Marco Garcia et al 2015, Ibrahim et al 2021). Our data is consistent with the idea that in the absence of retinal waves such transition is compromised. Layer 1 INs in β2-cKO mice have increased visual responses, suggesting increased thalamocortical inputs, and they receive reduced cholinergic inputs, suggestive of reduced long-range connectivity from neuromodulatory brain regions. Changes in connectivity onto input layer PYR and layer 1 INs could result from at least three different scenarios: (1) too many thalamocortical synapses are initially produced in the absence of cholinergic retinal waves, and/or (2) the elimination of excess thalamocortical synaptic inputs during thalamocortical circuit refinement does not occur normally in the absence of retinal waves, and/or (3) the transition from long range thalamocortical connectivity towards cortico-cortical and cholinergic connectivity is reduced. In follow-up studies, it will be important to further distinguish between those possibilities.

Growing circuits are likely to take advantage of all forms of activity-dependent plasticity available during their development. In the visual system retinal waves provide a plasticity mechanism for the visual cortex to develop basic properties, such as retinotopy, prior to eye opening. Accordingly, our data shows that β2-cKO mice have abnormal elevation maps consistent with the retinotopic preference of posterior V1 (Cang et al 2005). However, we show that other features of visual tuning, including orientation tuning, remain grossly intact in mutants. Such a result indicates that, in line with previous studies, some aspects of visual function may either be hardwired by activity-independent genetic processes (Bragg-Gonzalo et al 2021) or influenced by either sensory or other types of spontaneous activity than cholinergic retinal waves.

We also found that spontaneous activity manipulation during the first postnatal week led to later impairments in higher-order functions of the visual cortex, particularly behavioral state-dependent effects on visual physiology. Previous work by us and others has shown that the V1 response to sensory stimuli is modulated by an animal’s behavioral state, which allows animals to adjust sensory perception to changing behavioral demands (McGinley et al 2015, Saleem et al 2013, Stringer et al 2019, Vinck et al 2015). The circuit mechanisms through which behavioral states modulate sensory responses in V1 are still not well understood, but previous studies indicate that interneurons in layer 1 (Abs et al 2018, Cohen-Kashi Malina et al 2021, Ibrahim et al 2016, Roth et al 2016, Shlosberg et al 2006), and the development of thalamocortical connectivity is necessary for some aspects of state-dependent function (Murata & Colonnese 2018). Here we show that cholinergic retinal waves are critical for the development of behavioral state-dependent gain of visual responses of excitatory and layer 1 inhibitory circuits, suggesting the existence of a critical period for the establishment of behavioral state-dependent visual function that occurs prior to the onset of visual experience. Further, the period of cholinergic retinal waves overlaps with a vulnerable window for neurodevelopmental disorders with these disorders often manifesting deficits in state dependent brain function (Allene & Cossart 2010, Danka Mohammed & Khalil 2020, Kroon et al 2019, Llorca & Deogracias 2022, Marin 2012, Wong & Marin 2019). While spontaneous patterned activity and its role in early development remain poorly understood, our results shed light on the function of this highly conserved developmental process. Specifically, it suggests that prior to external sensory input, spontaneous activity plays a critical role in shaping high-order brain function. Understanding the precise role of these spontaneous waves on cortical development will require further work and a better understanding of how this activity shapes connectivity in the cortex.

## Materials and methods

### Animals

All mice were handled and maintained according to the regulations of the Institutional Animal Care and Use Committee at Albert Einstein College of Medicine and Harvard Medical School. Pax6α-Cre and β2fl/fl mouse lines were gifts of Michael C. Crair (Yale University, USA). C57/BL6 mice were used as wild-type (WT) unless otherwise specified. Pax6α-Cre::β2fl/fl (β2-cKO, Burbridge et al., 2014) mice were maintained on C57/BL6 background. Mice were bred on-site to generate the experimental groups.

### Single-nucleus RNA-sequencing

#### Widefield calcium imaging, and mapping wave+ and wave-regions in vivo

Mice were injected with AAV-GCaMP6f into one V1 region at P0-1 and headfixed while deeply anesthetized (3-4% isoflurane) with a surgical glue headcap over the skull at P7. Calcium imaging was done with LED stimulation through the skull after the mouse had been allowed to rise to a lighter plane of anesthesia (0.25%-0.5% isoflurane) to permit retinal wave activity (Ackman et al., 2012). Cortical patterned waves in V1 were imaged after mice were held at the lighter anesthetic state for the same amount of time as they were deeply anesthetized (to allow waves to reach a steady-state) and the entire V1 and part of surrounding V2 regions were imaged for about 20-30 minutes each. Based on the known regional retinal expression of Pax6α-Cre and V1 retinotopy that had largely been established by P6 in wild-type mice (Ackman et al., 2012, Fig. S1) wave frequency was measured in regions of interest starting in medial V1 and moving posterior towards the rear edge of V1. Decreases in wave frequency by region were noted and the ROI center of each region was marked on the gluecap over V1. The mouse was again deeply anesthetized and DiI was injected under each of the marked regions on the surface to denote the medial (wave+) and posterior (wave-) regions for micro-dissection (also see Supplemental movies S1 and S2 for examples of WT and cKO spontaneous cortical activity).

#### Tissue preparation, nuclear capture, and next-generation sequencing

Medial (wave+) and posterior (wave-) V1 tissue micro-dissected from β2-cKO and control mice (after imaging) was pooled such that tissue from between three and five mice constituted a given biological replicate. Three biological replicates were performed on different days to control for batch effects. Tissue was homogenized in homogenization buffer (HB) containing .25 M sucrose and (in mM): 25 KCl, 5 MgCl2, 20 Tricine-KOH, pH 7.8. and 2.5% Igepal-630 (Sigma). HB also contained 1 mM DTT, 0.15 mM spermine, and 0.5 mM spermidine along with phosphatase and protease inhibitors (Roche). Homogenization was performed on ice. After douncing the tissue ∼25 times, the homogenate was layered on top of a 30%-40% iodixanol gradient and centrifuged at 10,000g for 18 minutes. Small volumes of Bovine Serum Albumin (BSA; Sigma) and RNAsin (Promega) were included in HB and iodixanol solutions, to preserve high quality mRNA. Nuclei were recovered from the iodixanol interface and individually captured within microfluidic droplets alongside barcoded hydrogels via the inDrops approach(Zilionis et al 2017) in the single-cell RNA-sequencing core at Harvard Medical School. After cell encapsulation, primers were released by UV exposure. Four libraries of ∼2,500 nuclei were generated for each sample. Indexed libraries were pooled and sequenced three – four times on a Next-Seq 500 (Illumina) with sequencing parameters as follows: Read 1, 54 cycles; Read 2, 21 cycles; Index 1, 8 cycles; Index 2, 8 cycles. Data across bioreplicates were merged for analysis.

#### Data processing and analysis

Reads were mapped against a custom transcriptome built from Ensemble GRCm38 genome and GRCm38.84 annotation using Bowtie 1.1.1, after filtering the annotation gtf file (gencode.v17.annotation.gtf filtered for feature_type = “gene”, gene_type = “protein_coding” and gene_status = “KNOWN”). After mapping, sequence reads were linked to individual captured molecules by unique molecular identifiers (UMIs). Default parameters were used unless stated, as previously described (Cheadle et al 2018, Macosko et al 2015).

#### Quality control, cell clustering, and cell type assignment

In rare cases, two nuclei instead of one are captured within a given microfluidic droplet. To remove such instances from the dataset, we applied the R package DoubletFinder (https://github.com/chris-mcginnis-ucsf/DoubletFinder)(McGinnis et al 2019) to predict and remove doublets from each sample separately. This step aided in ensuring that unexpected doublets do not skew the analysis and the results. Roughly 6% of the nuclei were removed from the dataset using this approach. Next, nuclei that contained fewer than 250 or more than 3,000 UMIs were removed from the dataset as nuclei on the lower end of this spectrum may represent dead cells while nuclei on the higher end may represent surviving doublets. Cells containing greater than 5% mitochondrial read counts were also removed.

Further analysis of the dataset was performed using Seurat version 3. Data were log normalized and scaled to 10,000 transcripts per cell, and variable genes were identified using default parameters (Satija et al 2015). We limited the analysis to the top 30 principal components (PCs), and clustering resolution was set between 0.5 and 1.0. Clusters containing fewer than 100 cells were removed from the dataset as were clusters that appeared to express multiple distinct cell type markers. After assigning cells to general excitatory, inhibitory, and non-neuronal classes based upon the presence or absence of standard markers (e.g. Gad1 and -2 for interneurons, Neurod6 and Slc17a7 for excitatory cells, and the absence of either for non-neuronal cells), we separated out each general class and performed additional iterative clustering independently. This was effective in allowing us to filter out ambiguous cells and restrict the dataset to cells we could associate with a particular class with substantial confidence, and we detail the different classes we identified throughout the text (Tasic et al 2016, Tasic et al 2018). The final dataset included 65,744 nuclei across all conditions and bioreplicates. Average read depths for neuronal populations were 1,911 UMIs and 1,071 unique genes per cell.

#### Differential gene expression analysis

In this experiment, we included wave+ and wave-regions defined by 2-photon imaging from β2-cKO mice to identify spontaneous activity-dependent gene programs in V1. However, because there are also likely to be regional differences in transcription based upon topography – particularly in visual cortex where retinotopy is preserved – we also analyzed corresponding anatomical regions in Cre-negative littermate mice. To compare gene programs across these different regions in mice of both genotypes, we used Monocle version 2 (https://github.com/cole-trapnell-lab/monocle2-rge-paper), an R package specialized for quantitative analysis of single-cell data (Qiu et al 2017). This computational method has been found to robustly uncover differentially expressed genes in a number of different contexts, particularly in snRNAseq datasets with limited read coverage (Cheadle et al 2018). Genes whose differential expression met a rigorous false discovery rate (FDR) of less than 5% (FDR < 0.05) were considered statistically significant. Finally, we explored functional aspects of the data by performing gene ontology (GO) analyses on differentially expressed genes based upon PANTHER term enrichment (Mi et al 2013).

#### Fluorescence in situ hybridization (RNAscope)

Differential expression of a subset of genes was validated by fluorescence in situ hybridization (FISH) using the RNAscope platform. We used multiplexed detection kit version 1 reagents and followed the manufacturer’s instructions to perform experiments in fresh-frozen tissue. Probes used in the study include cFos (316921), Cadm1 (492361), Cadm2 (556101-C2), Tshz2 (431061), and Igfbp5 (425731). After FISH, sections were mounted on superfrost plus slides (Fisher) in fluoromount-G plus DAPI (ThermoFisher) and imaged at 63X on a LSM 710 Zeiss confocal microscope.

#### Quantitative PCR

Anterior and posterior V1 tissue from β2-cKO mice was micro-dissected, flash frozen in liquid nitrogen, and stored at -80 C until use. On the day of the experiment, tissue was thawed in 1 mL Trizol (Ambion; Naugutuck, CT), homogenized using a motorized tissue homogenizer (ThermoFisher), then extracted using chloroform. RNA from each sample was isolated using the RNeasy Micro kit (Qiagen) according to the manufacturer’s instructions. RNA concentrations were determined via nanodrop (ND 1000; NanoDrop Technologies, inc.; Wilmington, DE), then equal concentrations of RNA from each sample were converted into cDNA via SuperScript III First-Strand Synthesis System (ThermoFisher). Genes were amplified using forward and reverse primers and detected using Power Up Sybr Green reagent in a Quant Studio 3 Real-Time PCR system (ThermoFisher). Ct values were calculated and relative express was quantified using GAPDH expression as a reference control. Three biological replicates (n = mice) were included.

### Synaptic Density Analysis

#### Perfusion and Immunohistochemistry

Methods generally follow protocols in Favuzzi et al., 2017, 2021. Animals (P8, P21-25) were deeply anesthetized with sodium pentobarbital by intraperitoneal injection and transcardially perfused with PBS followed by 4% paraformaldehyde (PFA) in PBS. Brains were dissected out of the skull, postfixed for 2 hours in 4% PFA at 4ºC, embedded in 4% agarose and sectioned at 50μm on a vibratome (Leica). Brain sections were incubated with 0.3% Triton X-100 in PBS at room temperature for 4×15 minutes and then blocked for 2 hours in blocking solution (0.3% Triton X-100, either 10% Normal Goat Serum and/or 10% Normal Donkey Serum). Sections were then incubated overnight at 4ºC in primary antibodies in 0.3% Triton X-100 and either 5% Normal Goat Serum or 5% Normal Donkey Serum. The next day sections were rinsed in PBS at room temperature 4×15 minutes and then incubated with secondary antibodies at room temperature on an oscillating shaker for 2 hours in 0.3% Triton X-100 and 5% of either Normal Goat or Normal Donkey Serum. They were then rinsed 4×15 minutes in PBS and incubated with DAPI before mounting with Fluoromount-G (Southern Biotechnology). Antibodies used included: mouse anti-Synaptotagmin-2 (1:1000, ZFIN #ZDB-ATB-081002-25), mouse anti-gephyrin (1:500, Synaptic Systems #147 011), rabbit anti-NeuN (1:500, Millipore #ABN78), guinea-pig anti-NeuN (1:500, Millipore #ABN90P), guinea-pig anti-VGlut2 (1:2000, Millipore #AB2251), guinea-pig anti-Vglut2 (1:1000, Synaptic Systems #135404), guinea-pig anti-Vglut1 (1:1000, Millipore #AB5905), rabbit anti-Homer 1b/c (1:500, Synaptic Systems #160023), mouse anti-parvalbumin (1:1000, Sigma #P-3088), rabbit anti-DsRed (1:500, Clontech #632496), chicken anti-GFP (1:1000, Aves Lab #1020), guinea-pig anti-parvalbumin (1:2000, Swant #GP72), mouse anti-GAD65 (1:500, Millipore #MAB351R).

#### Image acquisition and analysis

For all analyses other than for layer Gad65 puncta images were taken in Layer 4 of primary visual cortex in the posterior (Wave negative) region. Gad65 puncta images were taken in the same V1 posterior region but in Layer 1. Tissue samples were imaged on an upright Zeiss LSM 800 confocal using a 40x oil immersion objective (1.4 nA, 2.5x digital zoom, 1024×1024 pixels). For synapse analysis single planes were analyzed using a custom Fiji script (ImageJ) as described previously (Favuzzi et al., 2017, 2021). Processing of images included noise reduction and smoothing and single channel images were converted to RGB. For PV synapse analysis a color threshold was set automatically to identify the cell body in layer 4, the perimeter was measured automatically, and a binary masked image with the cell body only was created. For all analyses a watershed-based method was used for bouton segmentation so that boutons were separated based on local minima of pixel gray-scale values. For presynaptic or postsynaptic boutons a color threshold was selected to segment boutons as isolated puncta. Comparison between the original images and the mask guided the choice of the threshold value, with the same criteria applied for each channel to all images within the same experiment. The “Analyze Particles” (in which the minimum size for presynaptic and postsynaptic structures was 0.20 μm and 0.10 μm, respectively) and “Watershed” tools were applied and a mask was generated. A merged image was created from all masks, converted to an 8-bit image, and the overlap between presynaptic boutons, postsynaptic clusters (and, for PV analysis, for cell body) was automatically detected as particles with a size greater than 0.05 μm2 in the “analyze particles” tool.

### In vivo electrophysiology

#### Surgery

Male and female animals between postnatal day 17 and 20 were anesthetized using isoflurane anesthesia and their body temperature was maintained at 37C using a closed loop heating pad. Their scalps were cleaned with alternating swabs of betadine and alcohol and incisions were made to remove a flap of scalp. The skull was scraped and conditioned with a 3% hydrogen peroxide solution to remove connective tissue. A gold ground pin soldered to a 200um tungsten wire was implanted above the cerebellum and secured with Flow-It dental composite. Optibond universal was applied to the skull followed by Ortho-Jet dental cement leaving a window over primary visual cortex using the dental cement to secure a 3D printed headpost based on the RIVETS system (cite). Animals were given dexamethasone and meloxicam for edema prevention and pain relief respectively.

#### Treadmill Habituation

Animals were allowed to recover for 24 hours after surgery before handling. Animals were rapidly habituated to head fixation by repeated handling multiple times per hour. Animals were introduced to head fixation at increasing lengths, a typical progression would be 10 secs, 15 secs, 30 secs, 1 min, 5 mins, 30 mins, 1 hour. Animals were removed from head fixation if they struggled or vocalized excessively.

#### In vivo recordings and state monitoring

We used linear silicon probes to perform high-density recordings of local field potentials (LFPs) and firing activity across all cortical layers in a head-fixed, unanesthetized mouse at p25, while monitoring the arousal state of the animal. The behavioral/arousal state of the animal was monitored through facial movements, pupil diameters, and locomotion on a treadmill. Animals were recorded at P25. Visual responses were evoked by presenting visual stimuli consisting of 20 degree visual angle, circular, drifting gratings of varying orientations and contrasts, directions, temporal frequency and size. Visual stimuli lasted 2 seconds with 1.5 seconds between trials and each unique grating was presented for 100 repetitions in a randomized order. Visual stimulation was controlled using Pyschtoolbox and custom written Matlab code and data was acquired using an Intan RHD USB interface board and an FLIR Blackfly camera for electrophysiology and video data, respectively.

#### Data analysis

Data were analyzed using custom written Matlab code along with publicly available data analysis packages. Single unit spiking activity was defined using Kilosort2.0 and facial movement and pupil size were extracted using FaceMap. Movement and stillness periods correlate with changes in pupil, locomotion and facial movement and were defined based on facial movement as it was the most reliable signal across recording sessions.

### In vivo calcium imaging

#### Surgery

Animals between P7 and P8 were injected with an AAV-Dlx-GCaMP6f in posterior visual cortex and allowed to express. At P21, animals underwent surgery as described above. In addition, a 3mm craniotomy was made over V1 and a cranial window was implanted and sealed using cyanoacrylate glue. This cranial window consisted of one 5mm cover glass attached to two 3mm cover glasses (Anderman methods paper) to prevent bone regrowth.

Imaging: Animals were imaged on a customized Thorlabs Bergamo 2-photon microscope equipped with the state monitoring described above. Signals from state monitoring and microscope signals were aligned using TTL pulses acquired on an Intan RHD system.

#### Data analysis

Data were analyzed using custom written Matlab code along with publicly available data analysis packages. Motion artifacts and ROI extract was performed using Suite2p.

### In vitro electrophysiology

P13-P15 mice for NDNF+ cells recording and P20-P30 mice for excitatory neuron recording used for in vitro electrophysiology experiments. Mice were decapitated, and the brain was removed and immersed in ice-cold oxygenated (95% O2 / 5% CO2) sucrose cutting solution containing 87 mM NaCl, 2.5 mM KCl, 2 mM MgCl2, 1 mM CaCl2, 1.25 mM NaH2PO4, 26 mM NaHCO3, 10 mM glucose and 75 mM sucrose (pH 7.4). 300μm thick coronal slices were cut using a Leica VT 1200S vibratome through posterior lateral V1 (corresponding to the wave negative region as determined by wide-field imaging from a different cohort of animals). Slices were recovered in a holding chamber with artificial cerebrospinal fluid (aCSF) containing 124 mM NaCl, 20 mM Glucose, 3 mM KCl, 1.2 mM NaH2PO4, 26 mM NaHCO3, 2 mM CaCl2, 1 mM MgCl2 (pH 7.4) at 34 °C for 30 minutes and at room temperate for 1 hour prior to recordings. Slices were then transferred to an upright microscope (Zeiss) with IR-DIC optics. Cells were visualized using a 40x water immersion objective. Slices were perfused with aCSF in a recording chamber at 2 mL/min at 30C. All slice preparation and recording solutions were oxygenated with carbogen gas (95% O2, 5% CO2, pH 7.4). Patch electrodes (3–6 MΩ) were pulled from borosilicate glass (1.5 mm OD, Harvard Apparatus). For all recordings patch pipettes were filled with an internal solution containing 130 K-Gluconate, 10 KCl, 10 HEPES, 0.2 EGTA, 4 MgATP, 0.3 NaGTP, 5 Phosphocreatine and 0.4% biocytin, equilibrated with KOH CO2 to a pH=7.3. All excitatory cells recording was done in the presence of 1um TTX. mEPSC and mIPSC were recorded in the presence of Gabazine(10um) and CNQX(20uM), AP5 (10uM) respectively. Cells were randomly patched based on the different layers they reside in. Recordings were performed using a Multiclamp 700B amplifier (Molecular Devices) and digitized using a Digidata 1440A and the Clampex 10 program suite (Molecular Devices). Voltage-clamp signals were filtered at 3 kHz and recorded with a sampling rate of 10 kHz. Recordings were performed at a holding potential of -70 mV. Cells were only accepted for analysis if the initial series resistance was less than 40 MΩ and did not change by more than 20% during the recording period. The series resistance was compensated at least ∼30% in voltage-clamp mode and no correction were made for the liquid junction potential.

### Data Analysis

Traces of synaptic events were downsampled and filtered with a Gaussian to reduce noise signals. The traces of the spontaneous excitatory events (EPSCs) was then performed using minianalysis software. Traces of synaptic events from excitatory cells were analyzed using clampfit software without any pre-processing. We calculated the amplitude and frequency of all events, as well as separated them into glutamatergic or cholinergic based on the kinetics of each event. Nicotinic cholinergic events have longer rise and decay times compared to AMPA mediated glutamatergic events. We used a rise time cut off window between 1.1ms and 5ms; and a decay time window between 3ms and 30ms. Cut off windows were inferred based on (Lamotte d’Incamps et al 2017).

### Statistical Analysis

No prior test for determining sample size was conducted. All statistical analyses were performed using Prism (GraphPad). Statistical significance was tested with non-paired, two-sided t-test, with a 95% confidence interval or One-Way ANOVA with Tukey’s correction for multiple comparisons. Data was tested for normality using the Shapiro Wilk test. If it did not pass normality, statistical significance was tested with Mann Whitney test. For some comparisons, two way ANOVA with Sidak multiple comparison test was also used. In some cases, One-Way ANOVA with Brown-Forsythe test was used in the cases where SD was not equal. All data are presented as mean + SEM unless otherwise stated.

### Cell tracing

Biocytin-filled layer 1 neurons were fixed in 4% paraformaldehyde and imaged on an upright Zeiss LSM 800 confocal in 100-200μm z-series with 0.5μm step size. Neurons were traced with the Simple Neurite Tracer tool (Fiji, ImageJ) and axons and dendrites were separated for length and Sholl analyses (Fiji). Statistical analysis was done in Prism (Graphpad) using an unpaired, two-sided t-test with a 95% confidence interval.

## Data availability statement

The snRNAseq data will be made available on the NCBI Gene Expression Omnibus (GEO) platform at the time of publication here: https://www.ncbi.nlm.nih.gov/geo/query/acc.cgi?acc=GSE211406. Reviewers can access the data prior to publication using the following access token: ibwtmkyuhzkzvap.

## Contribution statement

Initial conceptualization: T.B., L.C.; In vivo imaging in perinatal mice and model characterization; T.B., F.A.H., Single cell RNA-seq analysis: L.C., T.B.; Fluorescence in situ hybridization and qPCR: L.C.,U.V.; Perinatal Slice analysis: L.I., T.B.; Mature Slice analysis: D.D., T.B.; In vivo analysis of deficits in V1 in mature mice: J.R., R.B.B., L.S.; Writing, T.B., J.R., L.C., R.B.B. All authors contributed to editing. Supervision, G.F. and R.B.B; Funding Acquisition, G.F., R.B.B.

## Acknowledgements

Some schematics made using biorender.com. This work was supported by a NARSAD Young Investigator Award, a Simons Bridge to Independence Award, a Whitehall Award, and National Institutes of Health (NIH) grants R01EY034617, R21MH133097, R01EY034310 to R.B.B., by NINDS of the National Institutes of Health under F31NS120723 to J.R., by National Institutes of Health (NIH) grants R37MH071679 and R01NS081297 to G.F., by National Institutes of Health (NIH) grants R00MH120051, DP2MH132943, and R01NS131486 to L.C., a Simons Society of Fellows Junior Fellow award and Louis Perry Jones Postdoctoral Fellowship award to T.B.

---

**Supplemental Figure 1.**
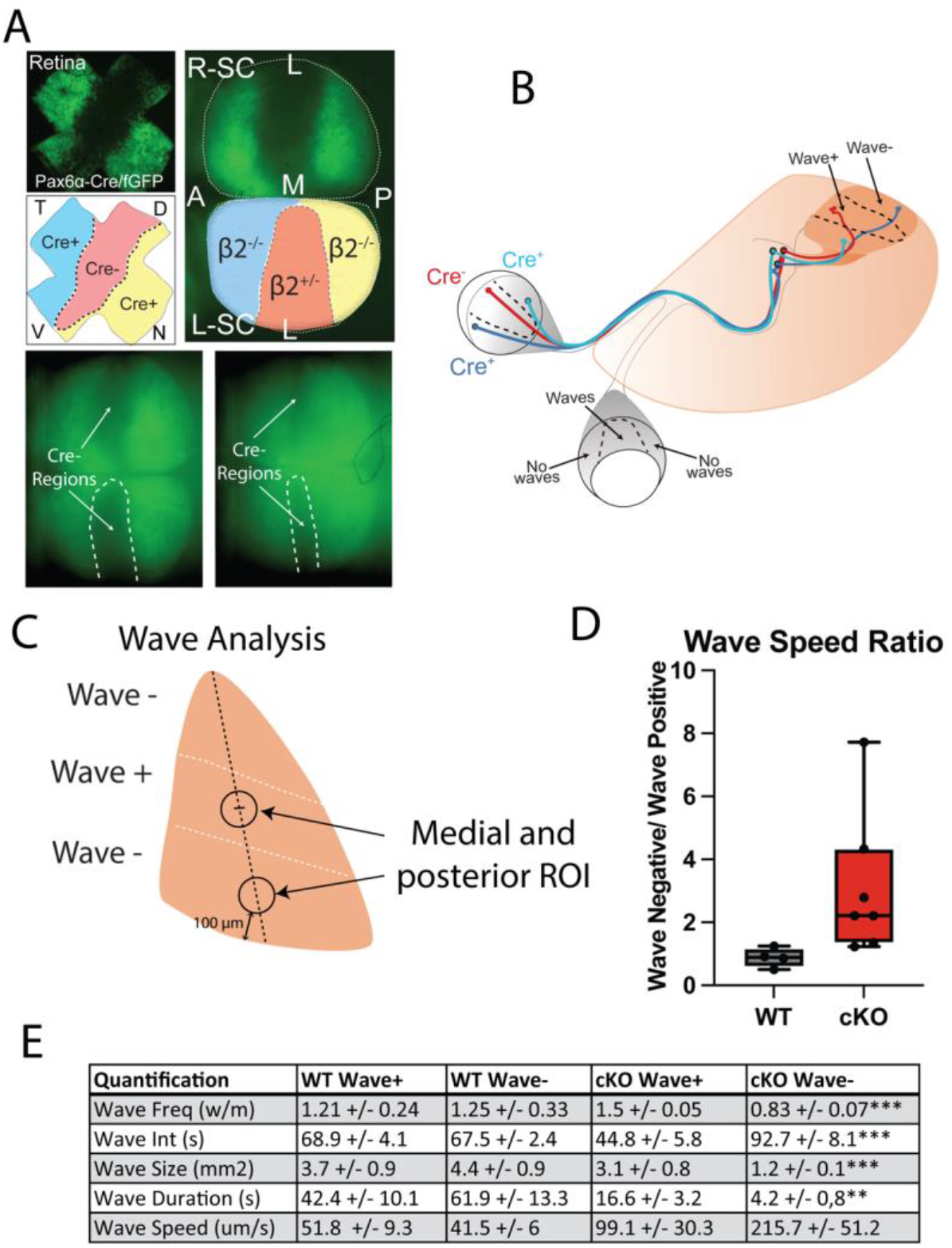
(Related to Figure 1). Pax6α-Cre::β2fl/fl mice specifically disrupts subregions of the retina with effects confined to corresponding subregions of V1. (A) Pax6α-Cre retinal expression shown with floxed eGFP (adapted from Burbridge et al., 2014). Cre expression varies between animals but Cre-region is always in the medial retinal in a dorsal to ventral strip. (B) Wave disruption method. Retinotopy is roughly set by P7 in V1, allowing approximation of Cre positive and negative regions (retinotopy shown in Fig. 5d, Ackman et al., 2012)(C) Analysis regions determined by retinotopy of Pax6α-Cre retinal regions and V1 retinotopy. (D) Wave speed analysis by region. E. Quantification of wave properties by region in WT and cKO mouse V1.

**Supplemental Figure 2.**
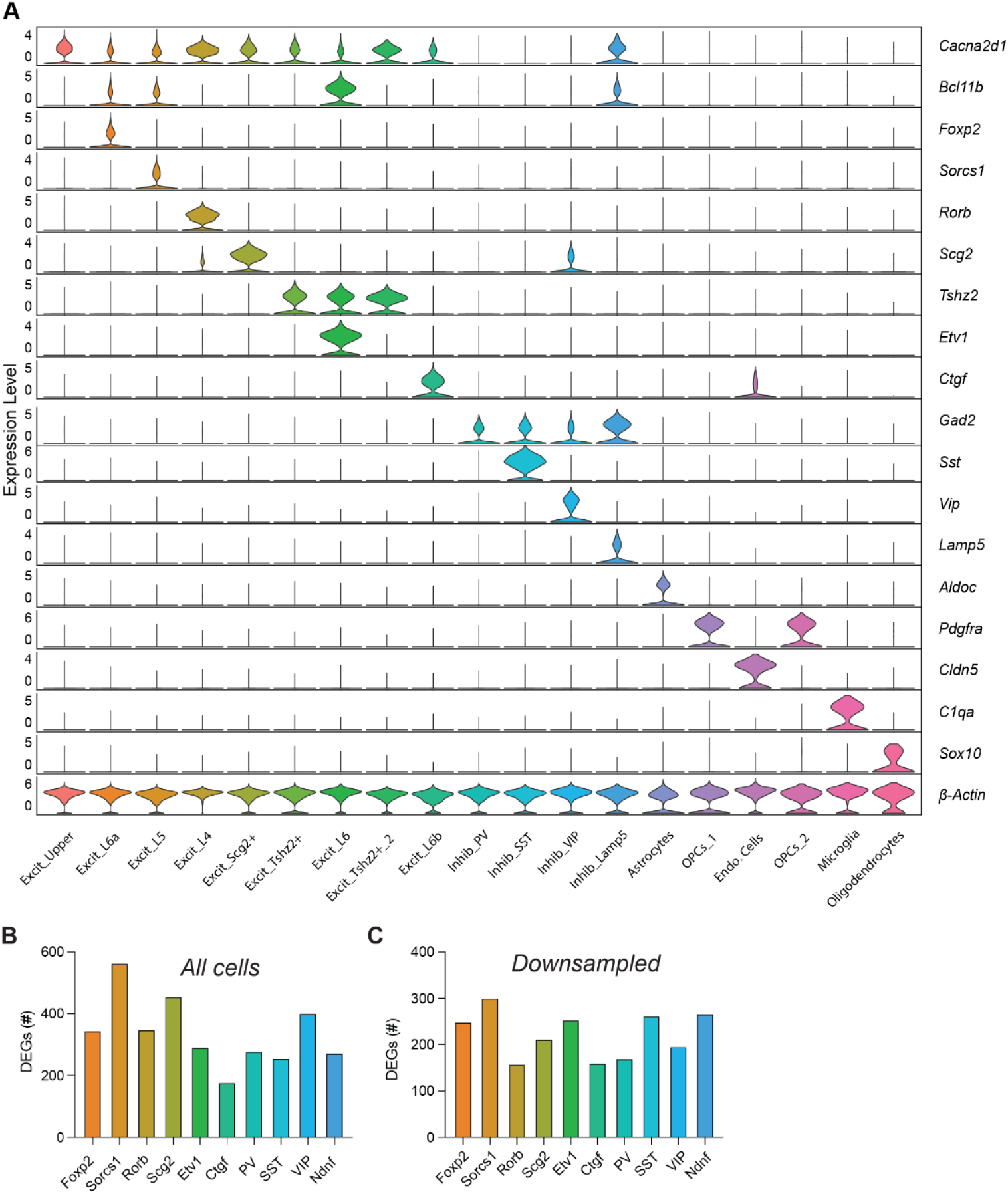
(Related to Figure 2). Identification of cell types in the snRNAseq dataset. (A) Violin plot demonstrating coherence between cell-type-specific marker genes (rows) and cell type identities (columns) in the snRNAseq dataset. Normalized expression level for each gene across all cell types shown on the left side of the Y axis. (B) The total number of differentially expressed genes (DEGs) in wave+ versus wave-regions of V1 in β2-cKO mice across neuronal subtypes. (C) Same analysis as in (B) but after down-sampling the number of cells in each cluster to 500.

**Supplemental Figure 3.**
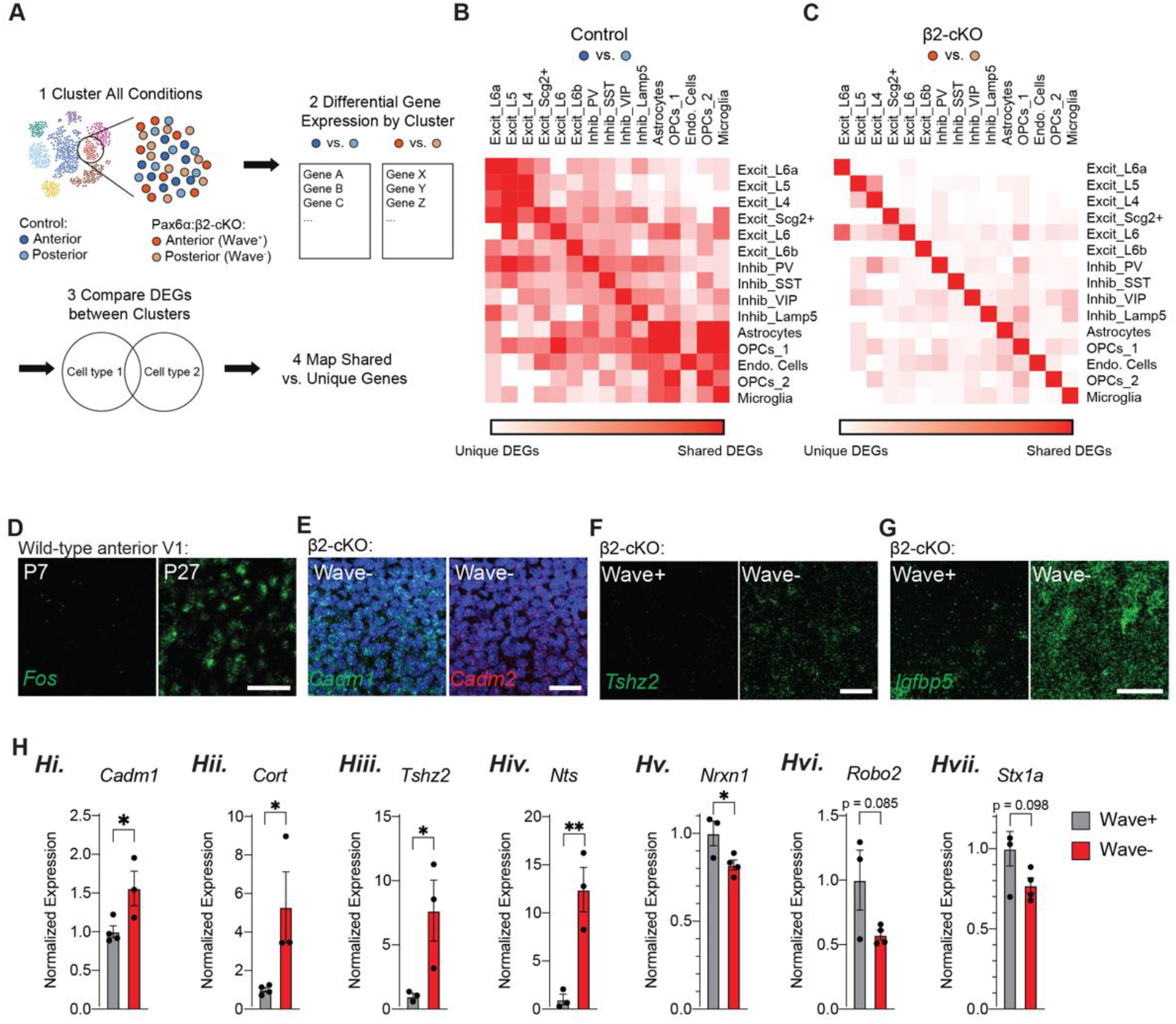
(Related to Figure 2). Characterization and validation of transcriptional changes in β2-cKO and control mice. (A) Schematic displaying the analysis pipeline to identify differentially expressed genes (DEGs) between cortical regions in all cell types in control (B) and β2-cKO (C) mice. (B) Heat map displaying overlap between DEGs, by cell cluster, when comparing anterior and posterior regions of V1 in control (Beta2fl/fl;Cre-negative) mice. White, region-specific genes in the two intersecting clusters are completely non-overlapping. Dark red, DEGs are identical between clusters. (C) Heat map displaying overlap between DEGs by cell cluster when comparing anterior (wave+) and posterior (wave-) regions of V1 in β2-cKO mice. Note that DEGs tend to be more cell-type-specific in the β2-cKO mice in which the level of activity differs between regions compared to control mice, where variability is primarily driven by topographical differences. (D) Confocal images demonstrating Fos mRNA expression in anterior V1 in wild type mice at P7 or P27, detected by fluorescence in situ hybridization (FISH). Scale bar, 50 m. (E) Expression of Cadm1 enriched in posterior V1 of β2-cKO mice compared to Cadm2. Scale bar, 50 m. (F) Expression of Tshz2 is upregulated in wave-negative regions of V1 in β2-cKO mice. Scale bar, 50 m. (G) Expression of Igfbp5 is upregulated in wave-negative regions of V1 in β2-cKO mice. Scale bar, 50 m. (H) Validation of transcriptional misregulation in wave-versus wave+ V1 in β2-cKO mice by qPCR: Hi. Cadm1; Hii. Cort; Hiii. Tshz2; Hiv. Nts; Hv. Nrxn1; Hvi. Robo2; and Hvii. Stx1a.

**Supplemental Figure 4.**
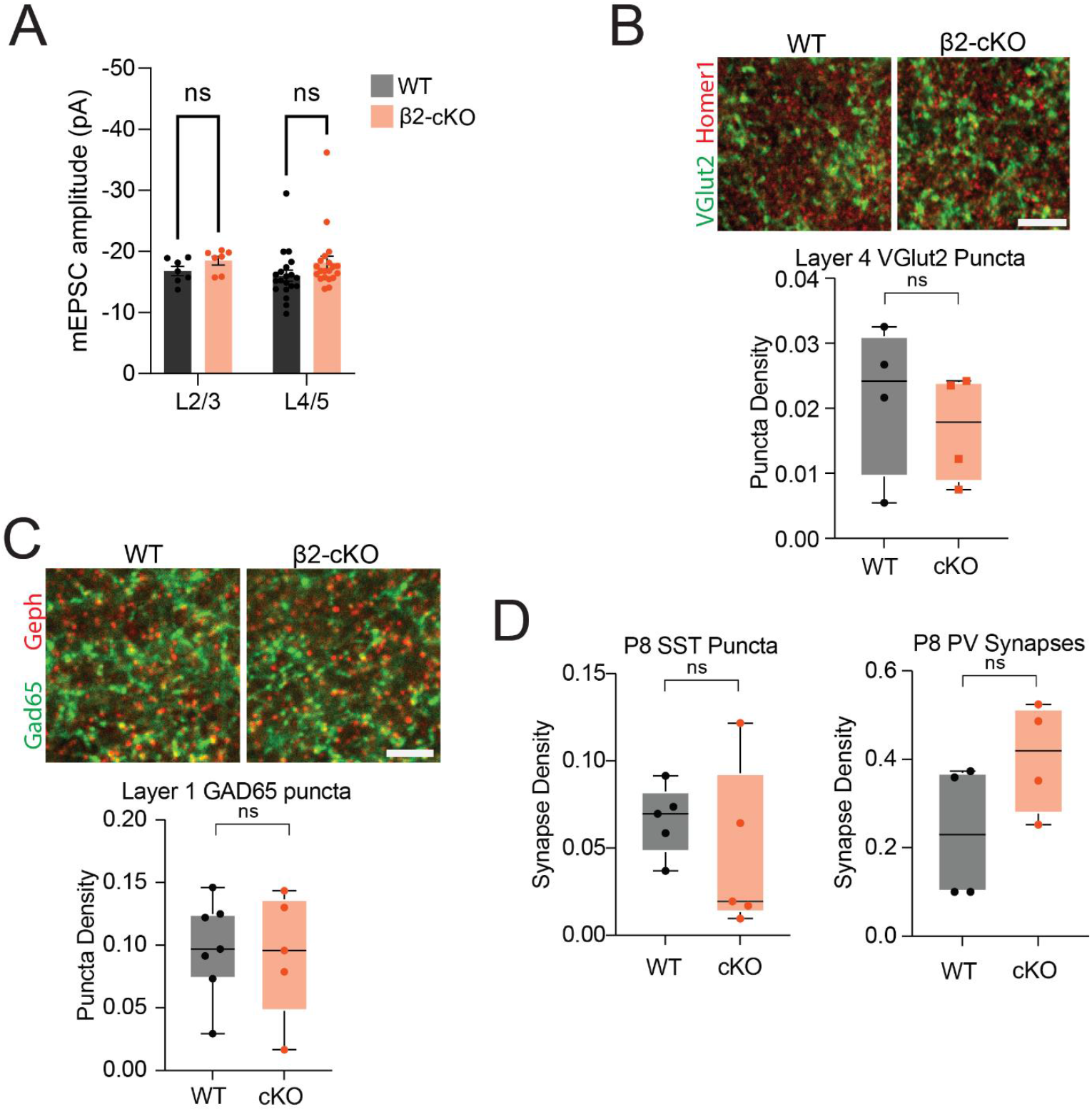
(Related to Figure 3): (A) mEPSC amplitude in pyramidal cells divided by layer. (B) Synaptic puncta analysis in P21 animals of layer 4 Vglut2 and (C) layer 1 Gad65 synapses measured in Layer 1. Here are no differences between control and mutant animals. N=4 control and mutants synaptic puncta. (D) Synaptic puncta analysis in P8 animals of L1 GAD65 puncta and PV puncta.

**Supplemental Figure 5:**
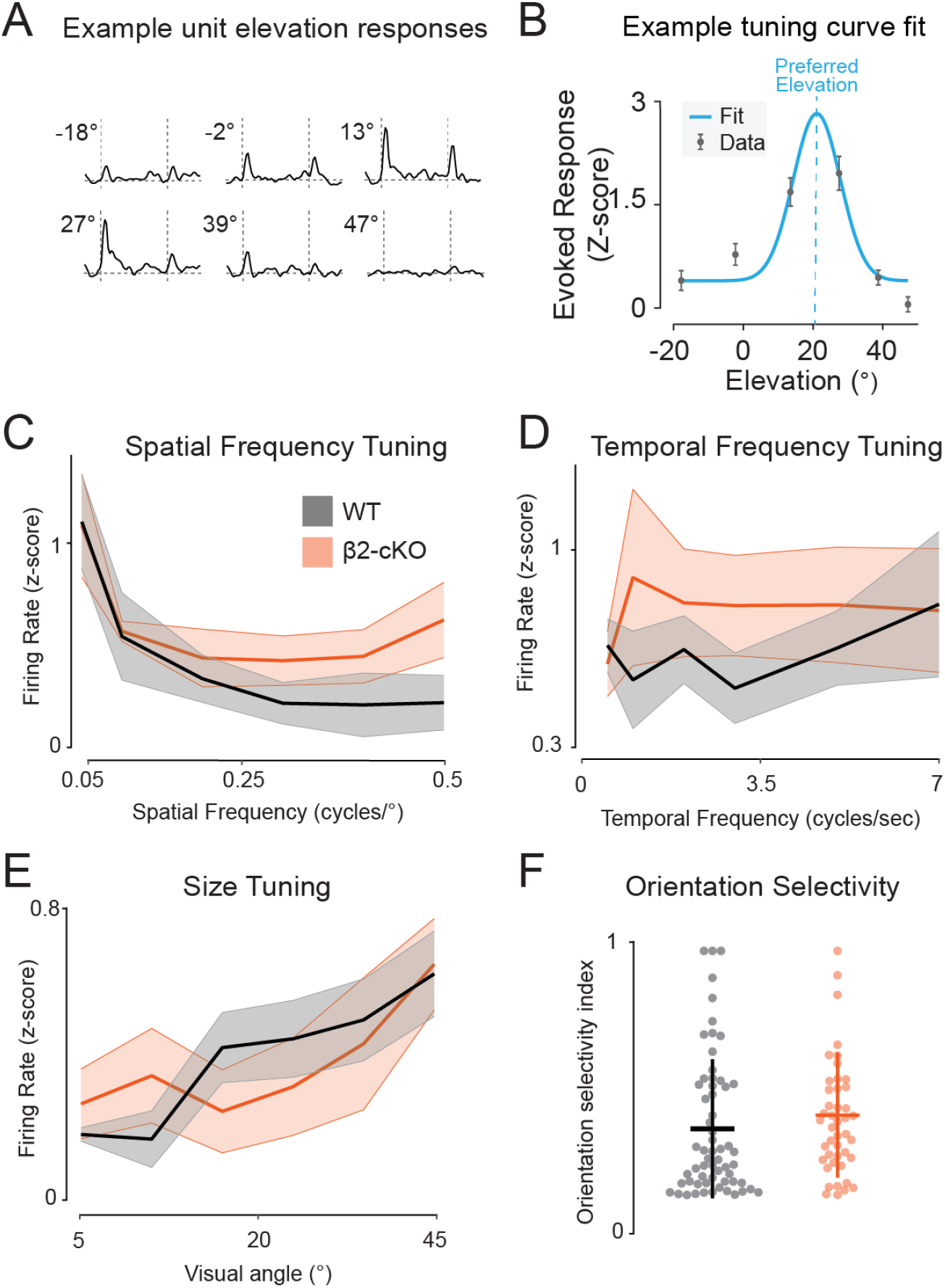
β2-cKO mice have some PYR receptive field properties intact. A) Example average responses to stimuli of different elevations. B) Gaussian tuning curve fit to data in A). C-E) Animal average spiking response to stimuli of varied C) spatial frequencies, D) temporal frequencies, and E) stimulus size. F) Orientation selectivity index of recorded units.

**Supplemental Figure 6:**
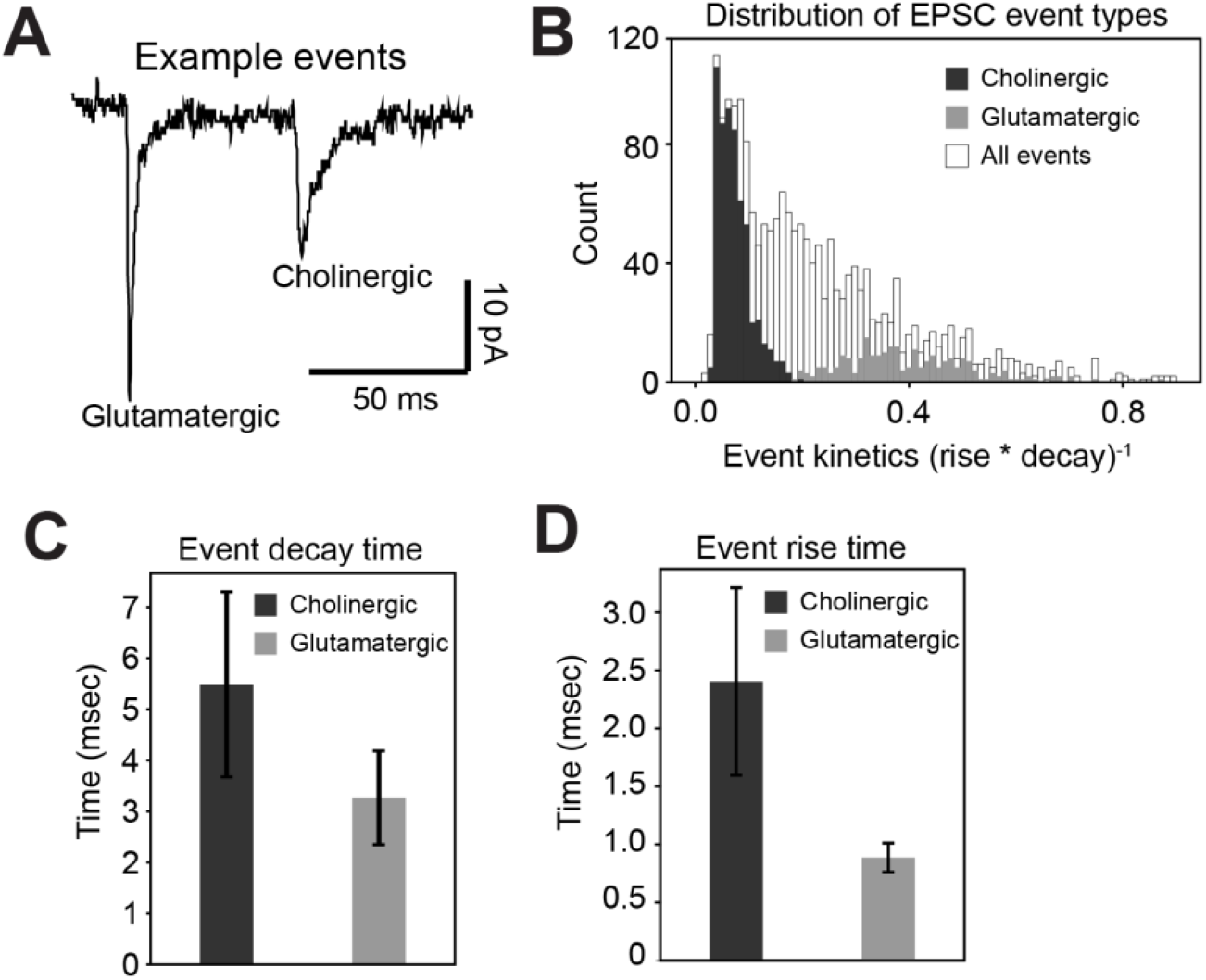
Separation of putative cholinergic and glutamatergic currents by kinetics. A) Example voltage clamp recording (clamped at -70mV) in a layer 1 cell. Left: faster glutamatergic current, right: slower cholinergic current. B) Histogram of kinetics of all events. C, D) Average rise time and decay time for all recorded currents. (error bars: s.d.)

**Supplemental Figure 7:**
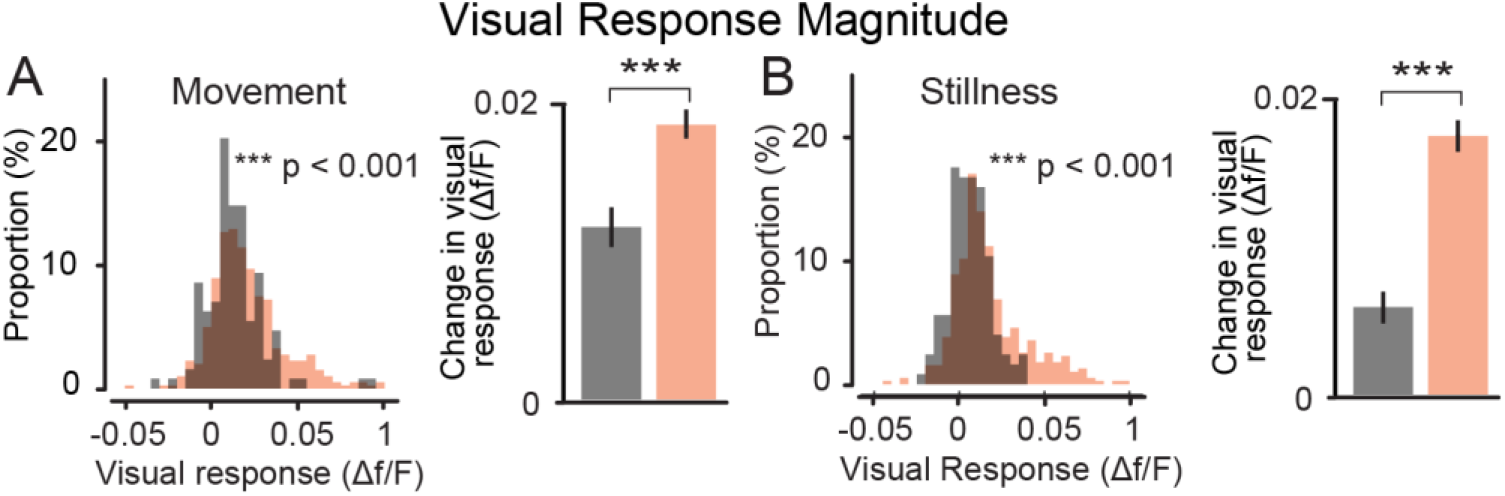
L1 IN visual responses are increased in β2-cKO mice across state. Histogram of average visual responses in Layer 1 INs during A. movement and B. stillness. (p < 0.001 Mann Witney U test) Left: histogram of all cells. Right: population mean and s.e.m.

**Supplemental Figure 8:**
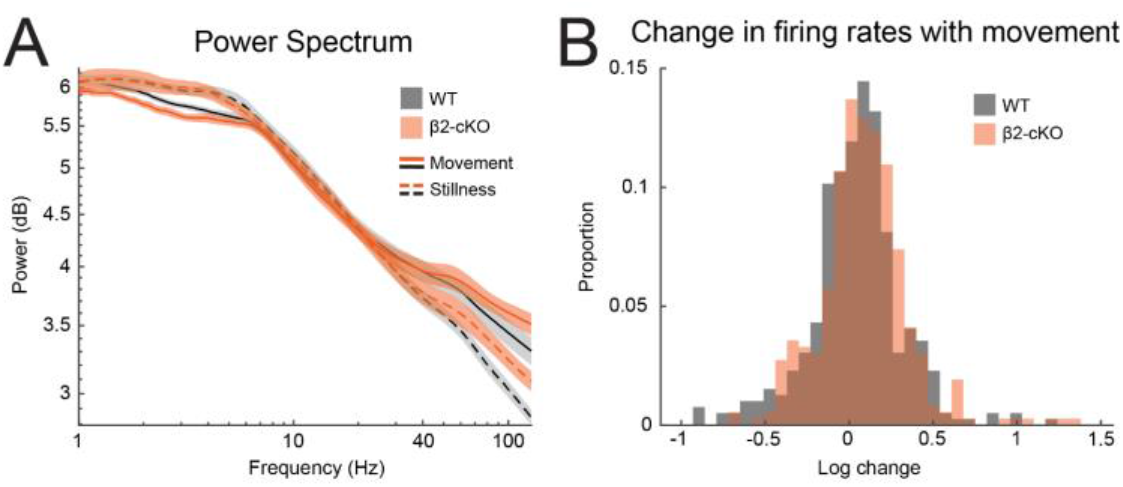
Power spectrum and firing rate modulation by arousal are similar in Beta2-cKO animals and controls. A) Power spectrum calculated for LFP recorded in layer 4. Calculated using wavelet transform of LFP data. B) Log change in firing rate from stillness to movement.

## References

Abs E, Poorthuis RB, Apelblat D, Muhammad K, Pardi MB, et al. 2018. Learning-Related Plasticity in Dendrite-Targeting Layer 1 Interneurons. Neuron 100: 684–99 e6

Ackman JB, Burbridge TJ, Crair MC. 2012. Retinal waves coordinate patterned activity throughout the developing visual system. Nature 490: 219–25

Allene C, Cossart R. 2010. Early NMDA receptor-driven waves of activity in the developing neocortex: physiological or pathological network oscillations? J Physiol 588: 83–91

Antonini A, Fagiolini M, Stryker MP. 1999. Anatomical correlates of functional plasticity in mouse visual cortex. J Neurosci 19: 4388–406

Antonini A, Stryker MP. 1996. Plasticity of geniculocortical afferents following brief or prolonged monocular occlusion in the cat. J Comp Neurol 369: 64–82

Ascoli GA, Alonso-Nanclares L, Anderson SA, Barrionuevo G, Benavides-Piccione R, et al. 2008. Petilla terminology: nomenclature of features of GABAergic interneurons of the cerebral cortex. Nat Rev Neurosci 9: 557–68

Bansal A, Singer JH, Hwang BJ, Xu W, Beaudet A, Feller MB. 2000. Mice lacking specific nicotinic acetylcholine receptor subunits exhibit dramatically altered spontaneous activity patterns and reveal a limited role for retinal waves in forming ON and OFF circuits in the inner retina. J Neurosci 20: 7672–81

Batista-Brito R, Fishell G. 2009. The developmental integration of cortical interneurons into a functional network. Curr Top Dev Biol 87: 81–118

Belousov AB, Fontes JD. 2013. Neuronal gap junctions: making and breaking connections during development and injury. Trends Neurosci 36: 227–36

Ben-Ari Y, Spitzer NC. 2004. Nature and nurture in brain development. Trends Neurosci 27: 361

Blankenship AG, Feller MB. 2010. Mechanisms underlying spontaneous patterned activity in developing neural circuits. Nat Rev Neurosci 11: 18–29

Bragg-Gonzalo L, De Leon Reyes NS, Nieto M. 2021. Genetic and activity dependent-mechanisms wiring the cortex: Two sides of the same coin. Semin Cell Dev Biol 118: 24–34

Bugeon S, Duffield J, Dipoppa M, Ritoux A, Prankerd I, et al. 2022. Atranscriptomic axis predicts state modulation of cortical interneurons. Nature 607: 330–38

Burbridge TJ, Xu HP, Ackman JB, Ge X, Zhang Y, et al. 2014. Visual circuit development requires patterned activity mediated by retinal acetylcholine receptors. Neuron 84: 1049–64

Cang J, Niell CM, Liu X, Pfeiffenberger C, Feldheim DA, Stryker MP. 2008. Selective disruption of one Cartesian axis of cortical maps and receptive fields by deficiency in ephrin-As and structured activity. Neuron 57: 511–23

Cang J, Renteria RC, Kaneko M, Liu X, Copenhagen DR, Stryker MP. 2005. Development of precise maps in visual cortex requires patterned spontaneous activity in the retina. Neuron 48: 797–809

Chandrasekaran AR, Plas DT, Gonzalez E, Crair MC. 2005. Evidence for an instructive role of retinal activity in retinotopic map refinement in the superior colliculus of the mouse. J Neurosci 25: 6929–38

Che A, Babij R, Iannone AF, Fetcho RN, Ferrer M, et al. 2018. Layer I Interneurons Sharpen Sensory Maps during Neonatal Development. Neuron 99: 98–116 e7

Cheadle L, Tzeng CP, Kalish BT, Harmin DA, Rivera S, et al. 2018. Visual Experience-Dependent Expression of Fn14 Is Required for Retinogeniculate Refinement. Neuron 99: 525–39 e10

Cohen-Kashi Malina K, Tsivourakis E, Kushinsky D, Apelblat D, Shtiglitz S, et al. 2021. NDNF interneurons in layer 1 gain-modulate whole cortical columns according to an animal’s behavioral state. Neuron 109: 2150–64 e5

Danka Mohammed CP, Khalil R. 2020. Postnatal Development of Visual Cortical Function in the Mammalian Brain. Front Syst Neurosci 14: 29

De Marco Garcia NV, Karayannis T, Fishell G. 2011. Neuronal activity is required for the development of specific cortical interneuron subtypes. Nature 472: 351–5

De Marco Garcia NV, Priya R, Tuncdemir SN, Fishell G, Karayannis T. 2015. Sensory inputs control the integration of neurogliaform interneurons into cortical circuits. Nat Neurosci 18: 393–401

Espinosa JS, Stryker MP. 2012. Development and plasticity of the primary visual cortex. Neuron 75: 230–49

Faust TE, Gunner G, Schafer DP. 2021. Mechanisms governing activity-dependent synaptic pruning in the developing mammalian CNS. Nat Rev Neurosci 22: 657–73

Fishell G, Kepecs A. 2019. Interneuron Types as Attractors and Controllers. Annu Rev Neurosci

Ford KJ, Felix AL, Feller MB. 2012. Cellular mechanisms underlying spatiotemporal features of cholinergic retinal waves. J Neurosci 32: 850–63

Ford KJ, Feller MB. 2012. Assembly and disassembly of a retinal cholinergic network. Vis Neurosci 29: 61–71

Fu Y, Tucciarone JM, Espinosa JS, Sheng N, Darcy DP, et al. 2014. A cortical circuit for gain control by behavioral state. Cell 156: 1139–52

Fukuda T, Kosaka T. 2000a. The dual network of GABAergic interneurons linked by both chemical and electrical synapses: a possible infrastructure of the cerebral cortex. Neurosci Res 38: 123–30

Fukuda T, Kosaka T. 2000b. Gap junctions linking the dendritic network of GABAergic interneurons in the hippocampus. J Neurosci 20: 1519–28

Galarreta M, Hestrin S. 2002. Electrical and chemical synapses among parvalbumin fast-spiking GABAergic interneurons in adult mouse neocortex. Proc Natl Acad Sci U S A 99: 12438–43

Grubb MS, Thompson ID. 2004. Visual response properties in the dorsal lateral geniculate nucleus of mice lacking the beta2 subunit of the nicotinic acetylcholine receptor. J Neurosci 24: 8459–69

Guillamon-Vivancos T, Anibal-Martinez M, Puche-Aroca L, Moreno-Bravo JA, Valdeolmillos M, et al. 2022. Input-dependent segregation of visual and somatosensory circuits in the mouse superior colliculus. Science 377: 845–50

Hensch TK. 2005. Critical period plasticity in local cortical circuits. Nat Rev Neurosci 6: 877–88

Hensch TK, Fagiolini M. 2005. Excitatory-inhibitory balance and critical period plasticity in developing visual cortex. Prog Brain Res 147: 115–24

Hrvatin S, Hochbaum DR, Nagy MA, Cicconet M, Robertson K, et al. 2018. Single-cell analysis of experience-dependent transcriptomic states in the mouse visual cortex. Nat Neurosci 21: 120–29

Huberman AD, Feller MB, Chapman B. 2008. Mechanisms underlying development of visual maps and receptive fields. Annu Rev Neurosci 31: 479–509

Ibrahim LA, Huang S, Fernandez-Otero M, Sherer M, Qiu Y, et al. 2021. Bottom-up inputs are required for establishment of top-down connectivity onto cortical layer 1 neurogliaform cells. Neuron 109: 3473–85 e5

Ibrahim LA, Mesik L, Ji XY, Fang Q, Li HF, et al. 2016. Cross-Modality Sharpening of Visual Cortical Processing through Layer-1-Mediated Inhibition and Disinhibition. Neuron 89: 1031–45

Jeon BB, Swain AD, Good JT, Chase SM, Kuhlman SJ. 2018. Feature selectivity is stable in primary visual cortex across a range of spatial frequencies. Sci Rep 8: 15288

Kirkby LA, Sack GS, Firl A, Feller MB. 2013. A role for correlated spontaneous activity in the assembly of neural circuits. Neuron 80: 1129–44

Kroon T, van Hugte E, van Linge L, Mansvelder HD, Meredith RM. 2019. Early postnatal development of pyramidal neurons across layers of the mouse medial prefrontal cortex. Sci Rep 9: 5037

Lamotte d’Incamps B, Bhumbra GS, Foster JD, Beato M, Ascher P. 2017. Segregation of glutamatergic and cholinergic transmission at the mixed motoneuron Renshaw cell synapse. Sci Rep 7: 4037

Lehrman EK, Wilton DK, Litvina EY, Welsh CA, Chang ST, et al. 2018. CD47 Protects Synapses from Excess Microglia-Mediated Pruning during Development. Neuron 100: 120–34 e6

Li Y, Fitzpatrick D, White LE. 2006. The development of direction selectivity in ferret visual cortex requires early visual experience. Nat Neurosci 9: 676–81

Llorca A, Deogracias R. 2022. Origin, Development, and Synaptogenesis of Cortical Interneurons. Front Neurosci 16: 929469

Macosko EZ, Basu A, Satija R, Nemesh J, Shekhar K, et al. 2015. Highly Parallel Genome-wide Expression Profiling of Individual Cells Using Nanoliter Droplets. Cell 161: 1202–14

Marin O. 2012. Interneuron dysfunction in psychiatric disorders. Nat Rev Neurosci 13: 107–20

Marquardt T, Ashery-Padan R, Andrejewski N, Scardigli R, Guillemot F, Gruss P. 2001. Pax6 is required for the multipotent state of retinal progenitor cells. Cell 105: 43–55

Marshel JH, Garrett ME, Nauhaus I, Callaway EM. 2011. Functional specialization of seven mouse visual cortical areas. Neuron 72: 1040–54

Martini FJ, Guillamon-Vivancos T, Moreno-Juan V, Valdeolmillos M, Lopez-Bendito G. 2021. Spontaneous activity in developing thalamic and cortical sensory networks. Neuron 109: 2519–34

Mazzaferro S, Bermudez I, Sine SM. 2017. alpha4beta2 Nicotinic Acetylcholine Receptors: RELATIONSHIPS BETWEEN SUBUNIT STOICHIOMETRY AND FUNCTION AT THE SINGLE CHANNEL LEVEL. J Biol Chem 292: 2729–40

McGinley MJ, Vinck M, Reimer J, Batista-Brito R, Zagha E, et al. 2015. Waking State: Rapid Variations Modulate Neural and Behavioral Responses. Neuron 87: 1143–61

McGinnis CS, Murrow LM, Gartner ZJ. 2019. DoubletFinder: Doublet Detection in Single-Cell RNA Sequencing Data Using Artificial Nearest Neighbors. Cell Syst 8: 329–37 e4

McLaughlin T, Torborg CL, Feller MB, O’Leary DD. 2003. Retinotopic map refinement requires spontaneous retinal waves during a brief critical period of development. Neuron 40: 1147–60

Mi H, Muruganujan A, Casagrande JT, Thomas PD. 2013. Large-scale gene function analysis with the PANTHER classification system. Nat Protoc 8: 1551–66

Monteiro P, Feng G. 2017. SHANK proteins: roles at the synapse and in autism spectrum disorder. Nat Rev Neurosci 18: 147–57

Mrsic-Flogel TD, Hofer SB, Creutzfeldt C, Cloez-Tayarani I, Changeux JP, et al. 2005. Altered map of visual space in the superior colliculus of mice lacking early retinal waves. J Neurosci 25: 6921–8

Murata Y, Colonnese MT. 2018. Thalamus Controls Development and Expression of Arousal States in Visual Cortex. J Neurosci 38: 8772–86

Niell CM, Stryker MP. 2010. Modulation of visual responses by behavioral state in mouse visual cortex. Neuron 65: 472–9

O’Leary DD, Crespo D, Fawcett JW, Cowan WM. 1986. The effect of intraocular tetrodotoxin on the postnatal reduction in the numbers of optic nerve axons in the rat. Brain Res 395: 96–103

Penn AA, Riquelme PA, Feller MB, Shatz CJ. 1998. Competition in retinogeniculate patterning driven by spontaneous activity. Science 279: 2108–12

Pereda AE. 2016. Developmental functions of electrical synapses. J Physiol 594: 2561–2

Pouchelon G, Dwivedi D, Bollmann Y, Agba CK, Xu Q, et al. 2021. The organization and development of cortical interneuron presynaptic circuits are area specific. Cell Rep 37: 109993

Qiu X, Mao Q, Tang Y, Wang L, Chawla R, et al. 2017. Reversed graph embedding resolves complex single-cell trajectories. Nat Methods

Reh RK, Dias BG, Nelson CA, 3rd, Kaufer D, Werker JF, et al. 2020. Critical period regulation across multiple timescales. Proc Natl Acad Sci U S A 117: 23242–51

Rochefort NL, Narushima M, Grienberger C, Marandi N, Hill DN, Konnerth A. 2011. Development of direction selectivity in mouse cortical neurons. Neuron 71: 425–32

Roth MM, Dahmen JC, Muir DR, Imhof F, Martini FJ, Hofer SB. 2016. Thalamic nuclei convey diverse contextual information to layer 1 of visual cortex. Nat Neurosci 19: 299–307

Saleem AB, Ayaz A, Jeffery KJ, Harris KD, Carandini M. 2013. Integration of visual motion and locomotion in mouse visual cortex. Nat Neurosci 16: 1864–9

Satija R, Farrell JA, Gennert D, Schier AF, Regev A. 2015. Spatial reconstruction of single-cell gene expression data. Nat Biotechnol 33: 495–502

Shah RD, Crair MC. 2008. Retinocollicular synapse maturation and plasticity are regulated by correlated retinal waves. J Neurosci 28: 292–303

Shatz CJ, Stryker MP. 1988. Prenatal tetrodotoxin infusion blocks segregation of retinogeniculate afferents. Science 242: 87–9

Shlosberg D, Amitai Y, Azouz R. 2006. Time-dependent, layer-specific modulation of sensory responses mediated by neocortical layer 1. J Neurophysiol 96: 3170–82

Sohl G, Maxeiner S, Willecke K. 2005. Expression and functions of neuronal gap junctions. Nat Rev Neurosci 6: 191–200

Stafford BK, Sher A, Litke AM, Feldheim DA. 2009. Spatial-temporal patterns of retinal waves underlying activity-dependent refinement of retinofugal projections. Neuron 64: 200–12

Stellwagen D, Shatz CJ. 2002. An instructive role for retinal waves in the development of retinogeniculate connectivity. Neuron 33: 357–67

Stiles J. 2011. Brain development and the nature versus nurture debate. Prog Brain Res 189: 3–22

Stringer C, Pachitariu M, Steinmetz N, Reddy CB, Carandini M, Harris KD. 2019. Spontaneous behaviors drive multidimensional, brainwide activity. Science 364: 255

Stuart T, Butler A, Hoffman P, Hafemeister C, Papalexi E, et al. 2019. Comprehensive Integration of Single-Cell Data. Cell 177: 1888–902 e21

Sun C, Speer CM, Wang GY, Chapman B, Chalupa LM. 2008a. Epibatidine application in vitro blocks retinal waves without silencing all retinal ganglion cell action potentials in developing retina of the mouse and ferret. J Neurophysiol 100: 3253–63

Sun C, Warland DK, Ballesteros JM, van der List D, Chalupa LM. 2008b. Retinal waves in mice lacking the beta2 subunit of the nicotinic acetylcholine receptor. Proc Natl Acad Sci U S A 105: 13638–43

Takesian AE, Bogart LJ, Lichtman JW, Hensch TK. 2018. Inhibitory circuit gating of auditory critical-period plasticity. Nat Neurosci 21: 218–27

Tasic B, Menon V, Nguyen TN, Kim TK, Jarsky T, et al. 2016. Adult mouse cortical cell taxonomy revealed by single cell transcriptomics. Nat Neurosci 19: 335–46

Tasic B, Yao Z, Graybuck LT, Smith KA, Nguyen TN, et al. 2018. Shared and distinct transcriptomic cell types across neocortical areas. Nature 563: 72–78

Triplett JW, Owens MT, Yamada J, Lemke G, Cang J, et al. 2009. Retinal input instructs alignment of visual topographic maps. Cell 139: 175–85

Trobiani L, Meringolo M, Diamanti T, Bourne Y, Marchot P, et al. 2020. The neuroligins and the synaptic pathway in Autism Spectrum Disorder. Neurosci Biobehav Rev 119: 37–51

Tuncdemir SN, Wamsley B, Stam FJ, Osakada F, Goulding M, et al. 2016. Early Somatostatin Interneuron Connectivity Mediates the Maturation of Deep Layer Cortical Circuits. Neuron 89: 521–35

Vinck M, Batista-Brito R, Knoblich U, Cardin JA. 2015. Arousal and locomotion make distinct contributions to cortical activity patterns and visual encoding. Neuron 86: 740–54

Wang L, Rangarajan KV, Lawhn-Heath CA, Sarnaik R, Wang BS, et al. 2009. Direction-specific disruption of subcortical visual behavior and receptive fields in mice lacking the beta2 subunit of nicotinic acetylcholine receptor. J Neurosci 29: 12909–18

Weaver IC. 2014. Integrating early life experience, gene expression, brain development, and emergent phenotypes: unraveling the thread of nature via nurture. Adv Genet 86: 277–307

Wong FK, Marin O. 2019. Developmental Cell Death in the Cerebral Cortex. Annu Rev Cell Dev Biol 35: 523–42

Yap EL, Greenberg ME. 2018. Activity-Regulated Transcription: Bridging the Gap between Neural Activity and Behavior. Neuron 100: 330–48

Zilionis R, Nainys J, Veres A, Savova V, Zemmour D, et al. 2017. Singlecell barcoding and sequencing using droplet microfluidics. Nat Protoc 12: 44–73

Zolnik TA, Connors BW. 2016. Electrical synapses and the development of inhibitory circuits in the thalamus. J Physiol 594: 2579–92

